# Transcriptional Variabilities in Human hiPSC-derived Cardiomyocytes: All Genes Are Not Equal and Their Robustness May Foretell Donor’s Disease Susceptibility

**DOI:** 10.1101/2024.04.18.584138

**Authors:** C. Charles Gu, Andrea Matter, Amy Turner, Praful Aggarwal, Wei Yang, Xiao Sun, Steven C. Hunt, Cora E. Lewis, Donna K Arnett, Blake Anson, Steve Kattman, Ulrich Broeckel

## Abstract

Human induced pluripotent stem cells (hiPSCs) are frequently used to study disease-associated variations. We characterized transcriptional variability from a hiPSC-derived cardiomyocyte (hiPSC-CM) study of left ventricular hypertrophy (LVH) using donor samples from the HyperGEN study. Multiple hiPSC-CM differentiations over reprogramming events (iPSC generation) across 7 donors were used to assess variabilities from reprogramming, differentiation, and donor LVH status. Variability arising from pathological alterations was assessed using a cardiac stimulant applied to the hiPSC-CMs to trigger hypertrophic responses. We found that for most genes (73.3%∼85.5%), technical variability was smaller than biological variability. Further, we identified and characterized lists of *“noise” genes* showing greater technical variability and *“signal” genes* showing greater biological variability. Together, they support a “genetic robustness” hypothesis of disease-modeling whereby cellular response to *relevant* stimuli in hiPSC-derived somatic cells from *diseased donors* tends to show *more* transcriptional variability. Our findings suggest that hiPSC-CMs can provide a valid model for cardiac hypertrophy and distinguish between technical and disease-relevant transcriptional changes.

## INTRODUCTION

Advances in human somatic cell reprogramming by selected transcription factors (Takahashi and Yamanaka, 2006; Yu et al., 2007) can erase epigenetic cell memory with reversion to a pluripotent state that is functionally similar to that of human embryonic stem cells (Guenther et al., 2010). The human induced pluripotent stem cells (hiPSCs) can be further differentiated into progeny cell types such as cardiomyocytes (CMs) (Braam et al., 2010; Kehat et al., 2001) and used as cell models for disease study (Dell’Era et al., 2015; Knollmann, 2013; Matsa et al., 2016a; Sinnecker et al., 2014; Tanaka et al., 2015). This technology provides a promising new platform and potentially unlimited cell source for regenerative medicine. As hiPSC-derived somatic cells become more accepted for disease modeling, the fidelity of the hiPSCs and their progeny cells becomes critical (Tapia and Schöler, 2016). To successfully apply the hiPSC-based model to studying human diseases, we must answer two essential questions. <u>First</u>, does the cell model faithfully depict the genetic impact on disease susceptibility in their donors? For example, when perturbed by external forces, e.g. drug stimulation, are there differences in cellular responses in the hiPSC-derived somatic cells between those generated from diseased individuals and those from others? <u>Second</u>, to what extent are the technical and undesirable variabilities due to reprogramming and differentiation processes *less than* the biological variabilities induced by the experimental conditions and donor’s genetic make-up?

While we know little about answers to the first question, there have been an increasing number of studies of clone to clone transcriptional variabilities in hiPSCs lines and their progeny somatic cells (Matsa et al., 2016b; Onder and Daley, 2012; Rouhani et al., 2014). Previous studies have investigated transcriptional variability in hiPSC lines, with the most recent studies showing substantial variability across hiPSC-derived cells (Germain and Testa, 2017; Matsa et al., 2016b; Onder and Daley, 2012). Multiple sources can contribute to the observed transcriptional variability in hiPSCs and progeny cells, such as processes used by hiPSC generation and somatic cell derivation, RNA-seq technique, controlled experiment conditions, and the donors’ individual biology. Some sources of technical variation are better understood than others. Numerous studies have been published about RNA-seq analysis showing that RNA-seq technical variabilities are generally smaller than biological variabilities, when using appropriate modeling (McIntyre et al., 2011). Other sources are less understood (e.g., transcriptional variabilities due to aberrations in hiPSC and progeny cell generation). Some factors may be improved upon (e.g., using later passages to erase epigenetic memory of cell origins), others are more intrinsically confounded and require deeper understanding and careful characterization (e.g., the expression of certain disease-associated genes also sensitive to hiPSC or progeny cells generation). One way to gain insight is to characterize and distinguish behaviors of genes under varying experimental conditions. For example, when stimulated by a potent drug, do some genes exhibit more variation in expression than others? Can the increased variabilities be explained solely by the stimulation or are they also influenced by the reprogramming process and/or donor disease status? Furthermore, are certain genes intrinsically variable and consequently less informative?

We address some of these questions using an experiment that utilizes highly pure populations of hiPSC-CMs (≥80% ventricular) derived in subjects selected from the HyperGEN-CiPS study (the Hypertension Genetic Epidemiology Network, a.k.a. HyperGEN, is a family-based cohort for studying genetic causes of hypertension and related diseases). For the analyses reported here, we have conducted 50 experiments across 25 clones derived from 18 hiPSC cell lines made from PMBCs of 6 individual HyperGEN subjects (“donors”) and one case of external sample of monogenic familial hypertrophic cardiomyopathy (FHM). To our knowledge, this is the first study to systematically characterize genome-wide transcriptional variabilities in *hiPSC-derived cardiomyocytes*, determine the relative source of variability, and their potential impact on disease modeling.

## RESULTS

### Generation of hiPSC lines and hiPSC-derived cardiomyocytes

The HyperGEN-CiPS study has generated hiPSC lines from blood cells collected from 248 HyperGEN subjects; the hiPSCs were differentiated to derive cardiomyocytes using proprietary protocols by FujiFilm Cellular Dynamics International (FCDI) as described in the METHODS section. For the purpose of this report, we selected 6 HyperGEN donors, made replicates of hiPSC lines through independent re-programming events, and from each hiPSC line generated up to three independent cardiomyocyte batches to characterize *inter-* and *intra-individual* variations. In total, this “clone experiment” dataset consists of 50 RNA-seq experiments performed on 21 *unique* hiPSC-CMs clones derived from 18 hiPSC lines created from blood cells in 6 HyperGEN subjects and 1 FHM sample provided by FCDI . Clinical characteristics of the donors are shown in Table 1. Repeated experiments were conducted in replicate samples at the levels of hiPSC generation, CM derivation, as well as drug stimulation & RNA-seq (Figure 1A).

**Table 1.** Clinical characteristics of study samples.

| SID | Sex | Race | Age | lvmht27 | lvmht27.2 | BMI | Diabetes |
| --- | --- | --- | --- | --- | --- | --- | --- |
| 1078 | Female | Black | 37 | 31.6 | 34.8 | 31 | Not Diabetic |
| 1082 | Female | Black | 37 | 23.7 | 24.9 | 22.1 | Not Diabetic |
| 1320 | Female | White | 60 | 39.2 | 48.3 | 37.6 | Not Diabetic |
| 1125 | Female | Black | 42 | 56 | 97.7 | 50.1 | Diabetic |
| 1135 | Female | Black | 43 | 46.9 | 61.4 | 33.5 | Not Diabetic |
| 1187 | Male | Black | 40 | 47.1 | 78.4 | 34.6 | Not Diabetic |
| 1178 | Monogenic familial hypertrophic myopathy (FHM)* |  |  |  |  |  |  |

\*No clinical data available

**Figure 1A.**
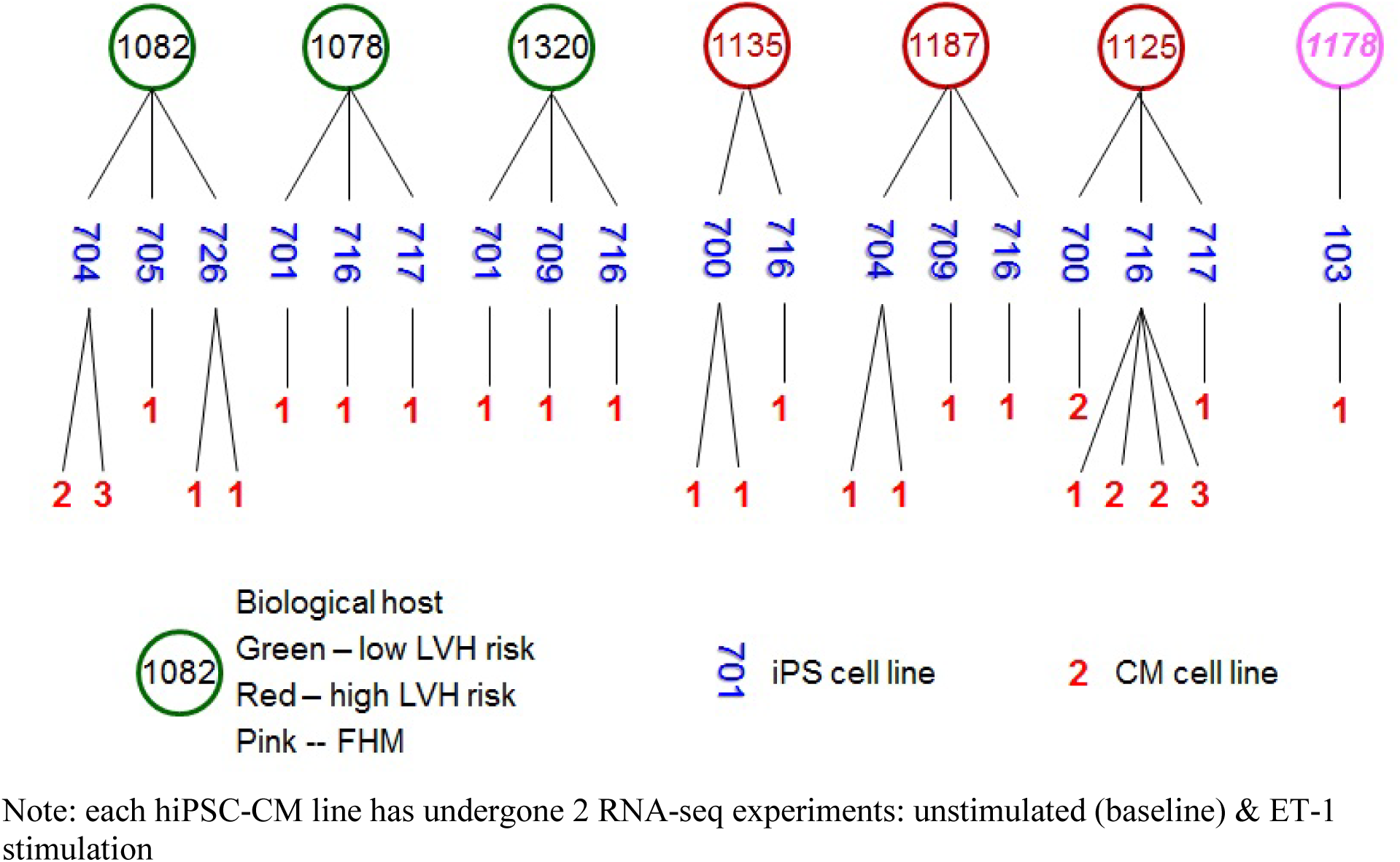
Design of the HyperGEN-CiPS clone experiment: Multiple replicates of iPSC lines & CM clones made from 6 HyperGEN donors: 3 LVH (circled in red); 3 normal (circled in green) subjects; plus 1 familial hypertrophic cardiomyopathy case (in pink). Blue numbers represent iPSC lines generated from the same donor, and red digit represent cardiomyocytes clones made from the same iPSC lines. There were 21 unique iPSC-CMs clones derived from 18 unique iPSC lines. Each CMs clone was divided into 2 aliquots for baseline and ET-1 stimulation, amounting to a total of 50 RNA-sequencing experiments.

**Figure 1B.**
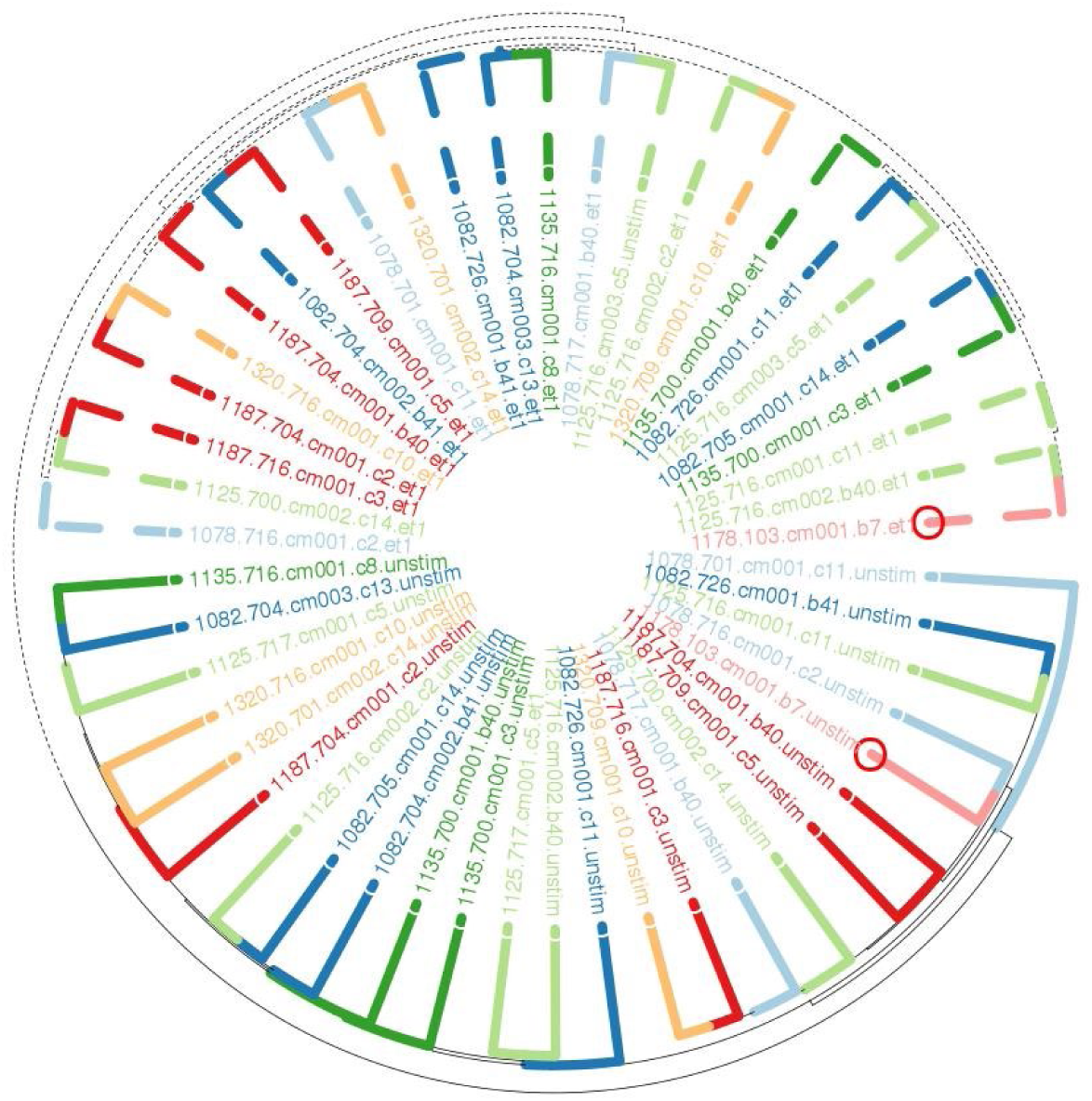
Recapture sample clustering: Hierarchical clustering was applied to expression levels normalized by voom in all transcripts passed QC. Clones obtained from the same donor were plotted in the same color. Small red circles indicate clones from the only FHM donor.

### Data pre-processing & quality control

For each hiPSC-CM line, a drug stimulation experiment was performed using endothelin-1 (ET-1) to induce molecular behavior mimicking LVH at the cellular level in cardiomyocytes. Genome-wide expression in both unstimulated (baseline) and stimulated cell lines were measured by RNA-seq (see METHODS). Quantified RNA-seq data were generated as counts data on a total of 48,737 transcripts. After initial QC and filtering by counts-per-million (cpm) a total of 16,791 transcripts remained for analysis. Examining for outlier experiments was performed by factor analysis implemented in the *RUVseq* package (Risso et al., 2014) that first removes “unwanted variations” in the data using empirical control genes identified by generalized log-linear model applied to the raw read counts, followed by visual examination of box plot of expression levels across all 50 experiments and PCA plot of the first 2 principal components extracted from the expression matrix. After removing the hidden unwanted variations, the log-ratios of read counts to the median across all samples were generally comparable and there was no experiment stand-out in the PCA plot. Closer examination of the QC box plots indicated that batch effect may have contributed to the hidden unwanted variation. We performed normalization on the counts data by *voom* (Law et al., 2014) implemented in *limma* (Ritchie et al., 2015; Smyth, 2005) using a linear model that includes batch, percentage of reads successfully mapped, and post plate counts (the number of cells that survive the plating process). Then the normalized expression residuals were used to check for sample clustering (Supplemental Figure 1B). Using all transcripts that passed QC, unbiased hierarchical clustering recaptured experiment conditions for all but two samples while clones from the same donors were grouped together most of the time with some hiPSC-CM lines from one donor occasionally clustering with those from other donors. More sophisticated modeling was applied to characterize and adjust for variations in the replicate experiments nested within either the same hiPSC-CM clones or the same hiPSC lines, and to separate “noise” genes from “signal” genes that can result in better clustering among other things.

### Proper modeling captures variabilities at different experimental stages

We applied generalized linear mixed effects models (GLMM) to properly capture sources of expression variabilities due to biological, experimental, and random factors by examining data from replicate experiments nested within the same donor, hiPSC clone, and CM line. Details of the models are given in Table S1 (#60&65). A series of additional nested models (#61∼64) were also fitted and compared with the null model by likelihood ratio test (LRT) to obtain a broadly inclusive list of “variable genes” (VR genes) due to any factors considered in the GLMM. The most restricted model (#65) included three fixed effects and five random effects, whereas the general (null) model (#60) included just the three fixed effects. To compare variance components at the different experiment stages for analyses described below, we fitted model #65 to the clone experiment data. Normalized expression levels were derived by adjusting all effects except for the group & individual effects estimated in the fitted model. Unless specified otherwise, further analyses reported below were based on the residuals as normalized expression levels.

### Biological effects are generally more variable than technical errors

For every gene/transcript that passed QC, we performed the GLMM analysis as described above to obtain estimates of variances for each source of variability modeled in #65. Except for 4 genes where the model did not converge, we obtained the variance component estimates for 16,793 genes. Comparisons of the various sources of variability are summarized in Table 2, in which “*Var(X)*” denotes the size of the variance component due to a given factor “*X*”. For example, *Var(tech)* denotes the size of variance due to the technical noise of RNA-seq. As shown, we examined the distributional properties of *Var(X)* for every factor *X* in model #65 including: technical noise (*tech*), hiPSC generation noise (*ips*), cardiomyocyte differentiation effect (*cms)*, individual donor background noise (*id*), and ET-1 stimulation experiment group (*grp*).

**Table 2.** Comparison of variance components of 5 random factors.

| Comparison of Variances | # of transcripts |
| --- | --- |
| **$\text{Var}(tch) + \text{Var}(cms) + \text{Var}(ips) \leq \text{Var}(ids) + \text{Var}(grp)$** | **10,096** |
| $\text{Var}(tch) \leq \text{Var}(cms)$ | 10,443 |
| $\text{Var}(tch) \leq \text{Var}(ips)$ | 11,645 |
| $\text{Var}(tch) \leq \text{Var}(ids)$ | 10,316 |
| $\text{Var}(tch) \leq \text{Var}(grp)$ | 13,661 |
| **$\text{Var}(tch) \leq \text{Var}(cms) + \text{Var}(ips)$** | **12,310** |
| $\text{Var}(cms) \leq \text{Var}(ips)$ | 11,243 |
| $\text{Var}(cms) \leq \text{Var}(ids)$ | 10,083 |
| $\text{Var}(cms) \leq \text{Var}(grp)$ | 12,931 |
| **$\text{Var}(cms) \leq \text{Var}(ids) + \text{Var}(grp)$** | **14,361** |
| $\text{Var}(ips) \leq \text{Var}(grp)$ | 11,876 |
| $\text{Var}(ips) \leq \text{Var}(ids)$ | 8,087 |
| **$\text{Var}(ips) \leq \text{Var}(ids) + \text{Var}(grp)$** | **13,524** |
| $\text{Var}(cms) + \text{Var}(ips) \leq \text{Var}(ids) + \text{Var}(grp)$ | 11,219 |
| $\text{Var}(ids) \leq \text{Var}(grp)$ | 12,670 |

*tch*: RNA-seq technical noise nested in CMs clone; *cms*: CMs differentiation effect nested within hiPSC lines; *ips*: hiPSC reprogramming effect nested in donor; *ids*: donor individual biological background; *grp*: ET-1 stimulation experimental effect.

The variation caused by ET-1 stimulation is of particular interest because it records systematic drug-induced response in cardiomyocytes and provides a model for susceptibility to the disease-state. The variability across individual donors reflects their variable biological backgrounds, parts of which are also relevant to the disease trait of interest. In contrast, the variations attributed to technical differences, and hiPSC and cardiomyocyte generation are undesirable noises. Therefore, for hiPSC-CMs to serve as a viable experimental model for studying LVH in donors, we would like to see that those variances due to noise (e.g., technical, hiPSC & CM generation) are far less than those due to functional and biological factors (e.g., cross-donor, drug stimulation, disease status, etc.).

As shown in Table 2, in the majority of genes (10,096 among 16,793), combined variances from technical and cell line generation (both hiPSC and hiPSC-CM) were less than that of donor individuality and experiment condition. Overall, it was more than 2-fold likely (11,219:5,574) that a randomly picked gene is more variable due to biology (donor variation or experiment conditions) than cloning.

Among the 3 noise factors, technical variance (RNA-seq) was smaller than variations due to CM (10,443) or hiPSC (11,645) generation, and less than the combined variations of the cell lines’ generation in a majority of genes (12,310, >73.3%). Transcriptional variabilities due to somatic cell generation (cms) were often less than those from hiPSC production (11,243, >66.9%), regardless of the size of variability due to ET-1 stimulation (also see Figure S1). For the biological factors, variations due to individual donor biology were generally smaller than that the stimulation (12,670, or >75.4% of genes tested), reflecting the impact of ET-1 as a potent cardiac stimulant.

Furthermore, comparing noise due to CM generation with that of other factors, it was generally less than that of stimulation (in 12,931 genes), and even more so than the combined effect of stimulation and individual donor biology in 14,361 genes (>85.5%). Similarly, the noise due to hiPSC generation was less than the variability by stimulation in 11,876 (70.7%) genes, and the combined effect of stimulation and individual biology in 13,524 genes (80.5%).

Interestingly, for randomly picked genes, the noise at baseline alone was equally likely to be caused by either hiPSC generation or the donor’s biological background. We surmounted this confounding issue partially by incorporating the perturbation paradigm (see panel (i) of Figure 2A and descriptions below). A complementary approach is to investigate which genes are more prone to noise factors arising from technical factors relating to producing the biological platform (e.g.hiPSC production, cellular differentiation, etc.). The data on replicate experiments gave us an opportunity to directly look into the behavior of such hiPSC-generation sensitive genes, as well as other groups of genes that might be sensitive to CM-generation or RNA-seq. We will describe these gene groups below in details.

**Figure 2A.**
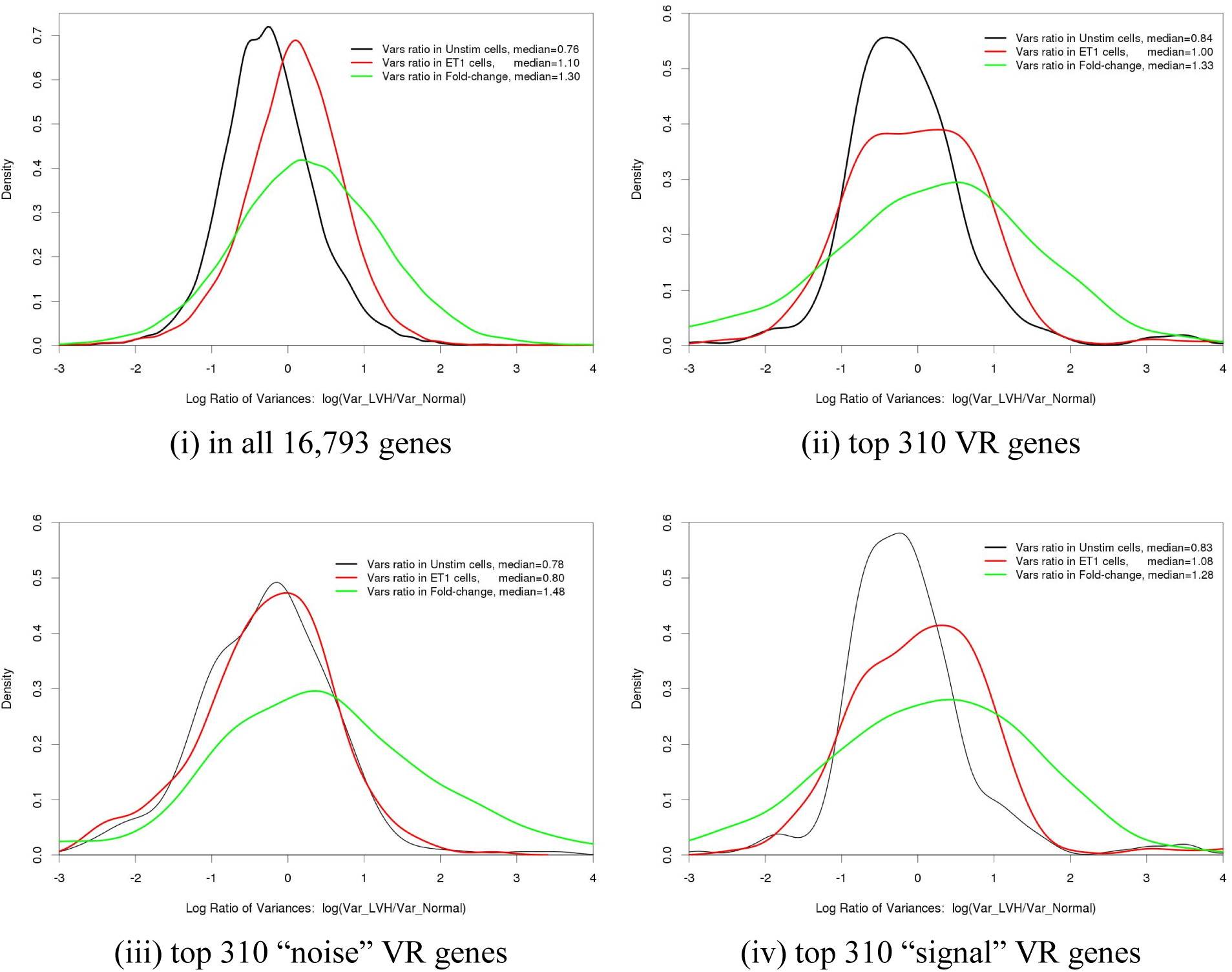
Comparison of distributions of LVH-to-Normal variance ratios in baseline versus ET-1 stimulated cell lines: Kernel density estimates of log-transformed ratios between variances in iPSC-CM clones from LVH donors and those from normal ones. The density estimates were calculated separately in baseline (unstim, colored in black)) and ET-1 stimulated cells (in red), as well as variance ratio of fold-changes (in green). Medians of the log-transformed ratios were displayed in the figure legends. Comparisons were done in (i) all genes; (ii) top 310 significantly variable (VR) genes as defined by LRT of GLMM models #65 (most restricted) and #60 (null); (iii) top 310 “noise” VR variable genes due to factors other than interesting biology defined by LRT of models #65 and #62; (iv) top 310 “signal” VR genes defined by LRT of models #62 and #60.

### The “genetic robustness” hypothesis of LVH

As mentioned previously, an important hypothesis about the validity of the hiPSC-based “disease-in-a-dish” model is that it recapitulates the donors’ disease genetics in terms of cellular response to external/internal perturbations. Our analysis of the clone experiment data showed that this “genetic robustness” hypothesis holds for the hiPSC-CM model for studying human LVH. This was done by genome-wide tests that compare expression variations in LVH- to non-LVH (‘normal-’) donor hiPSC-CMs, and across different experiment conditions.

More specifically, we performed transcriptome-wide tests to determine if the ratio of expression variances in LVH-donor hiPSC-CMs to that in normal-donor hiPSC-CMs is significantly greater than one. As shown in Figure 2A-(i), across all 16,793 analyzed transcripts, the distribution of LVH to normal variances ratio shifted to the right when changing from the unstimulated to the ET-1 stimulated condition. This was true no matter if we measured the expression levels separately in the LVH and normal hiPSC-CM groups (red) or as log of fold-changes (green). The same was also true in Figure 2A-(ii) when we limited analysis to the top 310 most variable genes based on LRT comparing the models #65 to #60. More interestingly, among the most variable genes, when measured separately in “noise” and “signal” genes (defined by differential expression (DE), see below), the shift was quite negligible among the “noise” VR genes as seen in Figure 2A-(iii), and more pronounced among “signal” VR genes shown in panel (iv) of Figure 2A. Furthermore, distributions of the variance ratios appear to be bimodal in panel (ii) & (iv) of Figure 2A, possibly reflecting the distinct behavior of the groups of genes responsive to stimulation and those more tied to a donor’s individual biology.

To further validate the robustness hypothesis, we performed differential gene expression analysis by re-fitting nested linear models on GLMM normalized expression values in three ways: donor LVH status was nested within stimulation group, stimulation was nested within donor LVH, and simply included donor-specific cardiomyocyte hypertrophy-by-stimulation interaction. After fitting the models, we counted significantly differentially expressed genes and compared the results under different conditions. As shown in Figure 2B, the resulting patterns corroborate the genetic robustness hypothesis. Specifically, when contrasting donor disease status (LVH vs normal individuals), there were more significant DE genes in ET-1 stimulated hiPSC-CMs than those in unstimulated ones; when contrasting the two experiment conditions (ET-1 stimulation versus unstimulated baseline), there were more DE genes in LVH-donor hiPSC-CMs than normal-donor hiPSC-CMs. As shown in Figure 2B-(i), when comparing the unstimulated group to the ET-1 stimulated group, the number of significant DE genes increased from 46 to 107 (26 overlapping; 20 & 81 unique to unstimulated & stimulated, respectively), and the increment was dominated by down regulated genes. In Figure 2B-(ii), switching from normal-donor to LVH-donor hiPSC-CMs clones, the number of significant DE genes by stimulation increased from 559 to 1,681 (448 overlapping, 111 & 1,233 unique to normal & LVH-donors, respectively), and the increment was dominated by up-regulated genes. In short, more genes are significantly variable in LVH-donor hiPSC-CM cells, possibly reflecting a less robust system leading to eventual manifest of LVH.

**Figure 2B.**
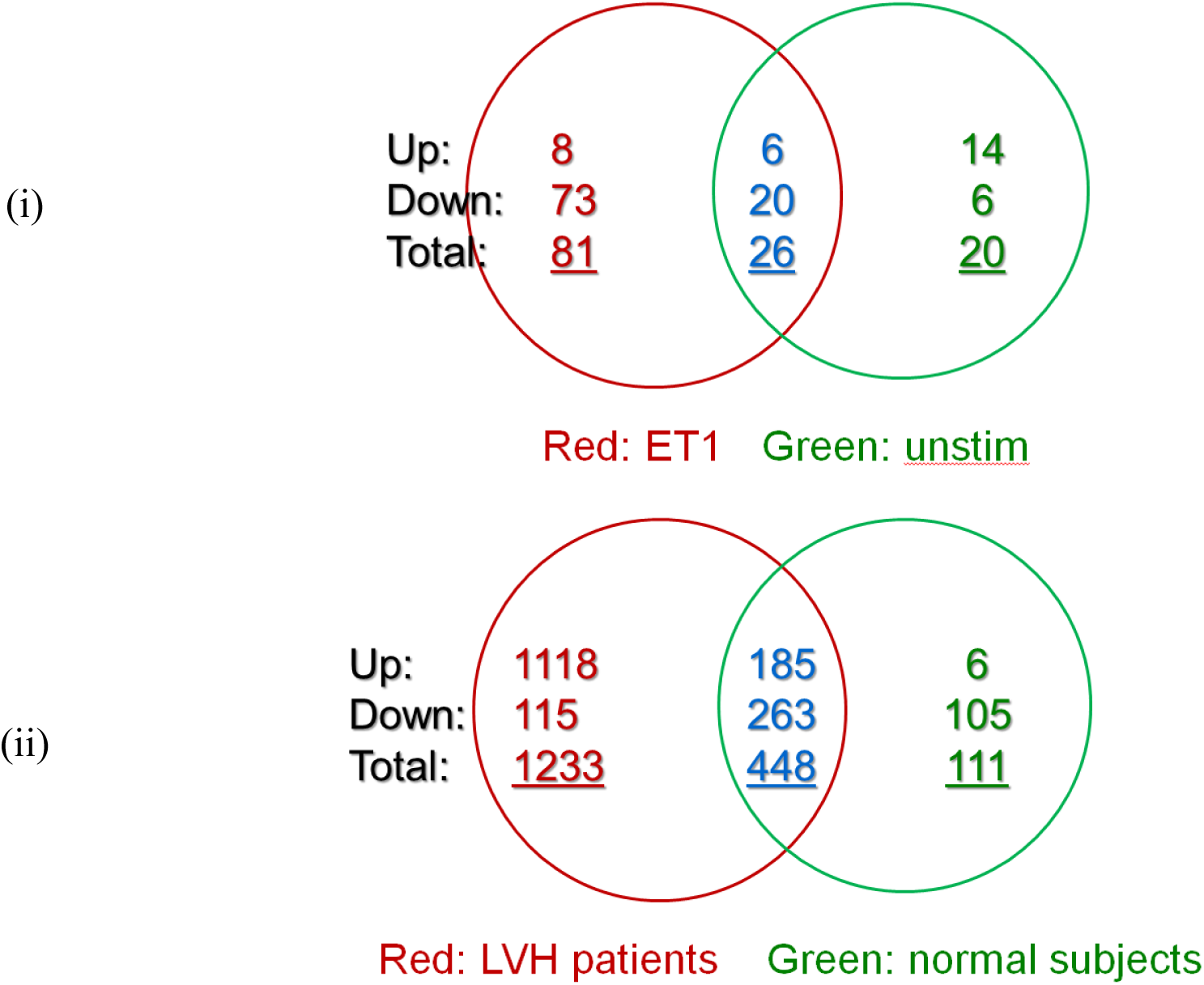
Compromised “genetic robustness” in iPSC-CMs from LVH donors: It is reflected by the increased numbers of significantly differentially expressed (DE) genes when: (i) comparing LVH to normal donors (i.e., DE genes due to donor LVH status), more genes were differentially expressed in ET-1 stimulated iPSC-CMs cells; (ii) comparing stimulated to baseline iPSC-CMs clones (i.e., DE genes responding to ET-1 stimulation), more genes were differentially expressed in cells made from LVH donors than in those from normal ones. The Venn diagrams were colored red (green) for the larger (smaller) sets of the DE genes, and included breakdown of numbers of up- and down-regulated DE genes.

### Existence of different types of genes sensitive to different steps of hiPSC cloning

Generating hiPSC-CMs is a cell reprogramming process that involves panels of transcription factors (OCT4, SOX2, NANOG, and LIN28), numerous participating target genes, and integration of drug resistance utilizing CM-specific promoter of the alpha-Myosin Heavy Chain (α-MHC).

Therefore, it is reasonable to hypothesize that the effects of reprogramming and subsequent differentiation on gene expression will vary by gene and stage of the process. That is, variation seen in expression analysis between hiPSC-CM lines from the same donor may have been influenced by random factors during hiPSC or hiPSC-CM generation. Some genes may be more sensitive to these random factors at different points in the process and thus contribute to technical noise rather than biological relevance.

If such “hiPSC-CM sensitive” genes were also involved in the etiology of the disease of interest, then their effects on the disease trait will be confounded in a study employing an hiPSC-CM based disease model. It is of critical importance that we learn as much as possible about which these sensitive genes are, and at which step(s) of the experiment their expression may fluctuate.

#### Sensitive Genes Defined by LRT

One may equate the types of sensitive genes to the groups of “variable genes” (VR) defined by likelihood ratio test (LRT) comparing 2 models in Table S1. For example, to use the hiPSC-CM model to study LVH we are mostly interested in separating genes whose expression levels are variable because of the donors’ LVH status, from those genes that show variability due to technical noises. Using a Bonferroni adjusted threshold of *α=*2.977*10^−06^ (=0.05/16793) and comparing model #65 to #62, we obtain a list of 62 VR genes that had significant non-zero effect on either of the three random factors for technical/RNA-seq noise, hiPSC or hiPSC-CM generation variations (i.e., “noise” genes). A similar LRT with 2 degrees of freedom comparing model #62 to #60 can be applied to obtain a list of 1,990 VR genes attributable to ET-1 stimulation or individual biology, both may be connected to donors’ disease status (i.e., differentially expressed, or DE, “signal” gene).

However, upon close examination of the variance components in the 62 genes estimated by model #65, variability for most of the genes on the list seemed to be attributable to hiPSC generation. This is seen in panel (i) of Figure 3A, where proportions of expression variance in the genes due to the 5 factors were summarized by the violin-plots as labeled. Except for a small number of genes, the largest fraction of variability was carried by the random effect due to hiPSC generation (reprogramming; median∼=0.629) and not hiPSC-CM generation (differentiation) or RNA-seq. Similar observations can be made for the list of “signal” DE genes obtained by LRT, *albeit* to a lesser extent. As seen in the violin plots in Figure 3A-(ii), while variabilities in most of the LRT-obtained DE genes were driven by ET-1 stimulation, only a very small fraction of them were attributable to donor individual biology. More importantly, a substantial proportion of the “signal” genes seemed confounded with (noise) variations due to hiPSC generation.

**Figure 3A.**
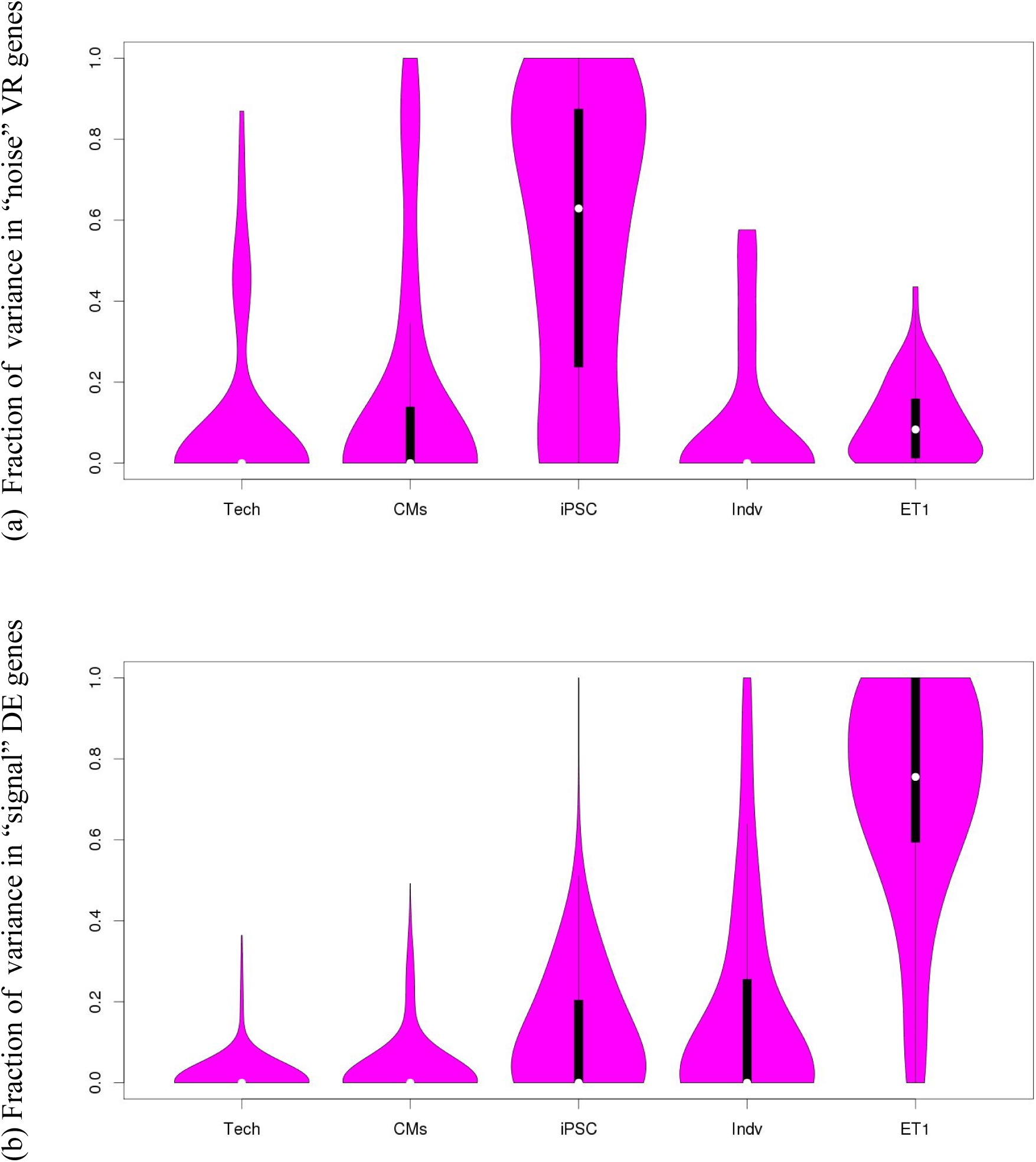
Distribution of proportions of variance in significant (α=0.05) variable (VR) genes selected by LRT: Using a significance threshold of α=0.05, we obtained 62 “noise” VR genes by contrasting GLMM model #65 to #62, and 1990 “signal” VR genes by testing model #62 against #60. Proportions of variance components due to each of the 5 random effects were estimated and displayed in a violin plot that combines a boxplot and a rotated kernel density plot on each side (see Methods). (i) For most of the 62 “noise” VR genes, major proportions of expression variances were attributable to the random effect of hiPSC generation; (ii) For most of the 1990 “signal” VR (DE) genes, proportions of expression variances were mostly driven by the effects of ET-1 stimulation.

**Figure 3B.**
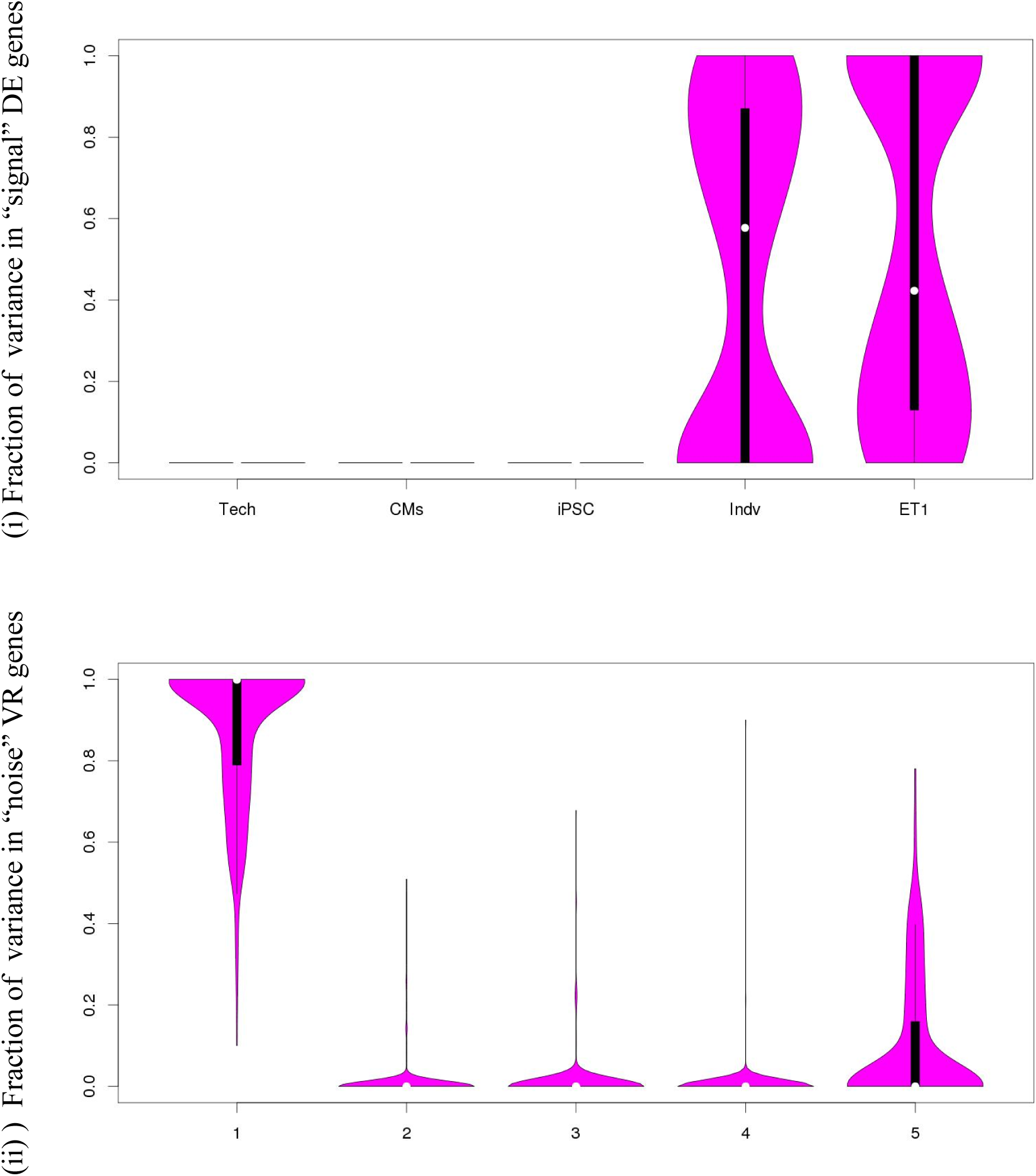
Distribution of proportions of variance in *refined* lists of VR genes selected by *top-and-bottom* quantiles of expression variances: Two examples were given here to show improved quality of the quantile-defined VR genes. (i) In the U list of VR (DE) genes comprised of union of G_3_ & G_4_ excluding those with the sum of variances due to the 3 “noise” sources below the 20% percentile, expression variances were exclusively attributable only to the effects of ET-1 stimulation and/or donor individual biology; (ii) In the “exclusive” list R_0_ of VR genes sensitive to just RNA-seq technical noise selected by top and bottom quantiles (see text), expression variance were largely carried by the technical fluctuation of RNA-seq.

Applying LRT in a stepwise fashion did not solve the problem: we obtained a sequence of gene lists for technical, hiPSC and hiPSC-CM generation and individual donor biology and stimulation that vary drastically in sizes (of 2, 1, 115, 253, and 2,024 genes). This can be due to multiple reasons: undesirable variations in the noise factors (e.g., RNA-seq experiments) may be confounded with other noises (e.g., hiPSC generation) or the disease biology (e.g., donor LVH status); it is well-known that estimate of degrees of freedom in mixed effects models can be unreliable, and the null distribution of LRT when comparing GLMM to GLM may be a mixture of chi-squares with different degrees of freedom. The latter may be addressed by using a more robust approach based on quantile as described below.

#### Sensitive Genes Defined by Quantile

We consider an alternative definition of VR genes by putting a threshold on the quantiles of their variance components estimated by the general model #65. Using a quantile threshold of top 1.8% (see Methods), we generate 5 types of VR genes, each due to a random effect in model #65, and analyze their behavior. Specifically, we will define the following 5 groups of variable genes: 3 groups due to noise and undesirable factors at each step of hiPSC generation, hiPSC-CM generation, and RNA-seq, respectively (called “noise” VR genes, or simply VR genes); and 2 groups due to donor individual biology and ET-1 stimulation (called “signal” DE genes, or simply DE genes). i.e., we apply the quantile threshold to the relevant variance components [Var(*type*) ≥ 98.2-percentile, where “*type*” may be “*tch*”, “*cms*”, “*ips*”, “*ids*”, and “*grp*”] to obtain ∼310 genes in each list and label them as: G_0_, G_1_, G_2_, G_3_, G_4_ for the lists of genes variable due to technical RNA-seq noise, hiPSC-CM generation, hiPSC generation, donor individual biology, and ET-1 stimulation. For example, G_0_ comprises of top 310 genes (largest technical variances) satisfying Var(*tch*) ≥ 98.2-percentile of technical variations, whereas G_4_ contains top 310 genes with Var(*grp*) ≥ 98.2-percentile of ET-1 induced variations.

As seen before, it is possible that for some genes multiple sources of variability are affecting their expression levels, although their relative contributions may vary. Therefore, we recognize that while the G lists defined above should capture the characteristic behavior of their respective (intended) source, the lists may not be exclusive (See Figure S2(a)). Indeed, as shown in Figure 4, where distribution of proportions of variance due to the 5 variance components were depicted by violin-plots (1 to 5) in each panel. There were quite a few genes in VR lists G_0_-G_4_ (meant to capture variability other than the stimulation) that also had non-negligible fractions of variance attributable to the stimulation (notice the non-zero medians in violin-plots #5 on panels (a)-(d) in Figure 4). This is probably because ET-1 is a highly potent cardiac stimulant. Nonetheless, carefully examining the violin-plots in Figure 4, we see that the overlaps were not large enough to amount to gross misrepresentation of the wrong sources.

**Figure 4.**
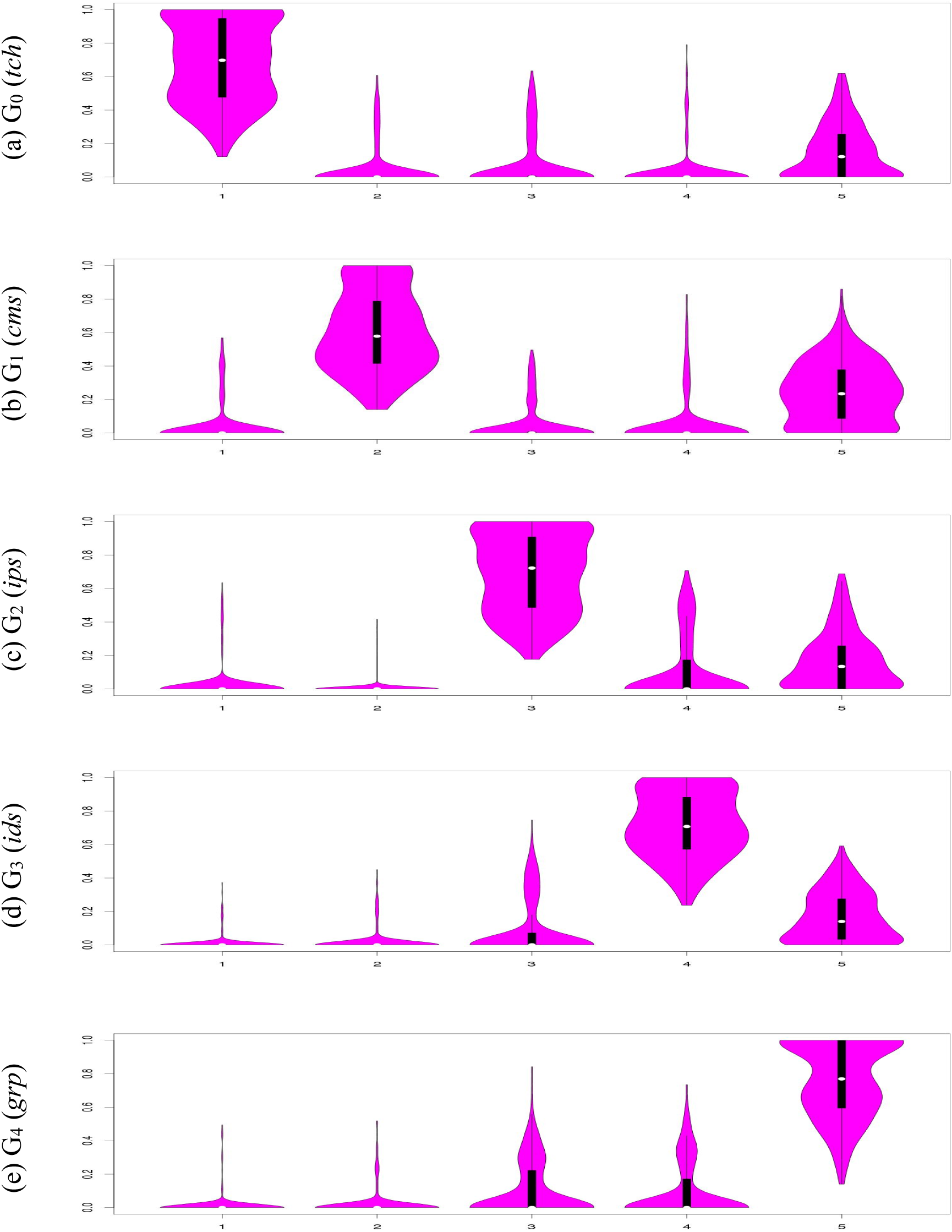
Distribution of proportion of variance in VR genes selected solely by top quantile (1.8%) of expression variances due to a specific factor: The VR gene lists G_0_ ∼ G_4_ were selected using a quantile threshold of 98.2% to obtain ∼310 genes (top 1.8%) most variable due to each of the five factors. Distribution of proportion of expression variances for each G list were shown by the violin plots, in panels (a) ∼ (e), respectively. In contrast to examples shown in Figure 3B, where the VR gene lists were obtained using 2 (top-and-bottom) quantile thresholds, expression variabilities in the G list of VR genes defined by a single quantile threshold tend to carry nontrivial fractions due some other factors. Abbreviations used: *tch:* RNA-seq technical noise nested in CMs clone; *cms:* CMs cloning effect nested within iPSC lines; *ips:* iPSC reprogramming effect nested in donor; *ids:* donor individual biological background; *grp:* ET-1 stimulation experimental effect.

#### Refined Lists of Sensitive Genes Defined by Quantile

The overlaps among the G lists defined above can be eliminated by refining their definitions on variance quantiles to gain further insights. For example, we may define the list “U” of jointly differentially expressed (DE) genes as the union of G_3_ & G_4_ subtract those genes for which the sum of variances due to the 3 “noise” sources below the 20% percentile (the threshold chosen to keep the size of U comparable to Gs). It is then expected that this group of genes should capture exclusively the “signal” expression changes due to ET-1 stimulation and/or the genetic background in the donors that include additional genetic effects related to their LVH status not affected by ET-1 stimulation. As seen in the violin-plots on panel (i) of Figure 3B, genes in the U list were indeed exclusively variable only because of stimulation or donor individuality, with no estimated variations attributable to the 3 noise factors. The property makes the U genes good candidates for improved clustering of our experiments. Indeed, hierarchical clustering based on the GLMM residuals of all U list genes resulted in near perfect grouping of the experiments by donors, and their nested hiPSC lines and CMs in an orderly fashion (Supplemental Figure S3; more clustering details in heatmap of pairwise distances in Figure S4).

Similarly, we can create 5 “exclusive” lists of VR genes that are sensitive to a single type of variability, by selecting genes with their variances in the top X-percentile with respect to a given type of variability AND the sum of their variances due to other sources in the bottom Y-percentile. For example, using such a pair of thresholds of top 10% and bottom 25%, we obtained 5 “exclusive” lists of VR genes R_0_, R_1_, R_2_, R_3_, and R_4_ that are exclusively sensitive to technical RNA-seq noise, hiPSC-CM generation, hiPSC generation, donor individual biology, and ET-1 stimulation, respectively. Unlike the G lists that all had ∼310 genes, the sizes of the R lists vary from 243 to 443 genes. Nonetheless, as expected, expression of genes in the R lists were mostly variable to a specific type of random effects. An example is shown by the violin-plots on panel (ii) of Figure 3B for genes in the list R_0_ where expression variabilities were largely carried by the technical noise of RNA-seq. The exclusive lists are also more informative. For example, using expression levels of VR genes in the exclusive list R_1_ which conveys sensitivity to hiPSC-CM generation, we can isolate the lone FHM sample from the HyperGEN donors (Supplementary Figure S5).

### Functional potential of genes in the stimulation responsive list G_4_

#### Cross validation by GWAS hits & eQTLs

To further understand the relationship between the variable genes and genetic variants underlying LVH we analyzed overlap between the VR gene lists and GWAS-detected SNPs associated with multiple LVH traits. To facilitate the comparison, we utilized information on the expression quantitative trait loci (eQTLs) that appear to regulate expression levels of some target gene(s). We considered whether any SNPs associated with two quantitative traits of LVH, left ventricular (LV) mass and ejection fraction (EF), might be affecting the disease development via regulating genes whose expression was also sensitive to LVH model induction via ET-1 stimulation. Specifically, do any of the GWAS-found SNPs regulate expression (in somatic cell types) of those G_4_ genes variable due to ET-1 stimulation?

We used data from the Genotype-Tissue Expression (GTEx) Project (V.6) that includes all significant eQTL SNPs that are potential regulatory variants of gene expression levels in 43 tissues/cell types based on genome-wide association analyses of expression measured by RNA-seq. For each gene in G_4_, we recorded all their eQTL SNPs detected at a significance level of α=1.0E-5 in any of the 4 LVH related tissues (left ventricle, atrial appendage, coronary artery, and aorta) analyzed by GTEx. These eQTL SNPs were checked against significant SNPs detected by GWAS analysis of LV mass or EF.

To be inclusive, a significance threshold of α=1.0E-3 was used to obtain a total of 2,080 SNPs (in 1,246 genes) from previous GWAS for LV mass and 1,648 (in 1,065 genes) for EF. Comparing the lists with that from the GTEx-derived eQTLs, we found 4 GWAS-detected SNPs that are eQTLs: 2 of them associated with LV mass and 2 with EF, all detected in African American (AA) samples (none in European American (EA) samples). Both SNPs associated with EF, rs4244842 and rs11228427, are located near the MRGPRD gene on Chromosome 11, about 15kb and 12kb downstream, respectively. Of the 2 SNPs associated with LV mass, rs309134 is located in an intron of DARS-AS1 on Chromosome 2, and rs636049 in an intron of MRPL21 on Chromosome 11. We note that the two genes on Chromosome 11, MRPL21 & MRGPRD, are 76kb apart on the same strand. There was a report in dbGaP about another SNP, rs10496739, in the DARS-AS1 gene associated with Echocardiography (Vasan et al., 2007). While the target gene of rs309134 is MCM6 on the complement strand of Chromosome 2, the target of all three Chromosome 11 eQTLs was RP11-757G1.6, a small ncRNA gene, whose functionality is relatively unknown.

A closer look at the expression levels in the GTEx data revealed that for each of the overlapping SNPs, the effect sizes of differential gene expression was substantial in the 4 LVH relevant cell types, as seen in Supplemental Figure S6. For rs309134, top 2 leading effects were in LV and aorta; for rs636049, expressions of its target gene were abundant in many cell types, and the eQTL has substantial positive effects in all 4 heart-related tissues. For the 2 SNPs overlapping with EF GWAS, the effects were limited to 1∼2 types cells, and one of the SNPs (rs11228427) had opposite effects in LV and aorta.

#### Enrichment of known pathways and gene-sets

Findings of overlapping GWAS-found SNPs with eQTL SNPs regulating those responsive to the ET-1 stimulation prompted us to ask if the overlap can also be seen at the pathway/gene set level. We verified this by enrichment analysis using information about known pathways and gene-sets collected by the Molecular Signatures Database (MSigDB; Subramanian et al., 2005). We tested for significant enrichment of the pathways in genes harboring the significant GWAS SNPs (“GWAS genes”) and in G_4_, and subsequently identifying shared gene-sets/pathways. For comparison, we also performed the same enrichment analyses in lists G_0_, G_1_, G_2_, and G_3_. Based on our selection procedure for the G lists, if the choice largely reflected true underlying variabilities, then we should see most overlap between GWAS gene list and the G_4_ list (responsive to ET-1 stimulated), with only a smaller overlap with the G_0_ list (technical noise).

As shown in Table 3-A, there were 1,246 unique genes harboring (2,080) SNPs that are associated with (log) LV mass (α=1.0E-3). Parsing through a total of 15,752 pathways and gene sets compiled by the MSigDB, we found 2,237 pathways enriched in the LV mass GWAS gene list. Among the 5 variable gene lists G_0_, …, G_4_ (with largest variance components due to sources from technical noise, CMs generation, etc., with equal number of genes selected), G_4_ has the largest overlap (30 genes out of 303) with the GWAS gene list while G_0_ & G_3_ (representing variabilities due to tech noise and individual biology) had the smallest overlap (14 & 9 genes respectively). At the pathway level, the two lists also had the smallest overlap of significantly enriched gene sets (215 & 1, respectively). However, G_1_ shared slightly more significantly enriched pathways with the GWAS gene list than G_4_ did (430 vs 412, respectively). We reasoned that some pathways sensitive to CM-generation may also be involved in the development of LVH or ET-1 stimulation. In other words, while the analysis separated out technical from biologically-relevant variabilities, the G designation for each group may contain genes that are variable due to multiple possible sources. To narrow down on genes that are more likely variable due to a unique source, we focus on the “exclusive” VR gene lists R_0_, R_1_, R_2_, R_3_, and R_4_ defined in previous section by selecting from the top 90 percentiles of variance of the corresponding random effects, while restricting that the sum of variances of the other 4 sources was in the bottom quartile. Here we further require an equal number of genes from the top of each list, to make 5 new lists T_0_ ∼ T_4_, each with 243 genes (See Figure S2(b)). To a large extent, genes from each of the T lists are variable in the clone experiment largely due to a single source of variability. Repeating the pathway analyses on the new T lists, we obtained the results shown in Table 3-B. Overlap of the T lists with the GWAS genes was not very different at the gene-level, with T4 having the most (17 genes) and T0 the least (7 genes). However, it was quite interesting that, although all lists had the same number of genes, there were large differences in terms of *significantly* enriched pathways across the lists. T_4_, variability driven largely by ET-1 stimulation, had the most (649), likely because ET-1 is a potent stimulant that affects many pathways. Many of the affected pathways were probably also relevant in the etiology of LVH, because it also had the most number (102) of enriched pathways overlapping with those enriched in the GWAS list. Similar to G3, T3 also had very few enriched pathways and none overlapped with those enriched in the GWAS gene list. For T1 & T2, variability due to CM differentiation and hiPSC generation, significantly enriched pathways were also few (none for T1 and 9 for T2). For T0, the largely “noise” gene list, there were quite a few (111) significantly enriched pathways. However, only a few (9) overlapped with those also enriched in the GWAS gene list. The overlapping gene sets with a size of 200 genes or less are given in Supplemental Table S2.

**Table 3-A.**
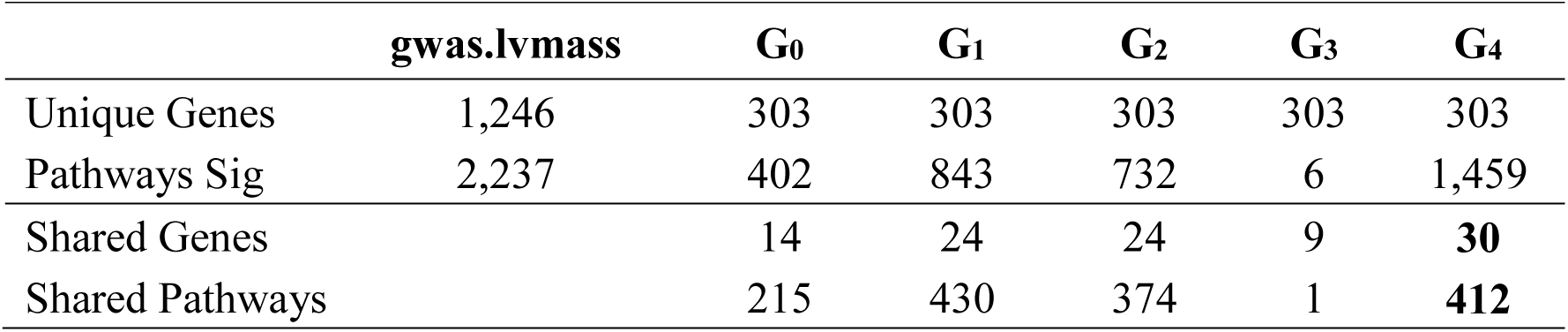
Overlap of pathways significantly enriched in GWAS-detected genes and those in the lists G_0_ ∼ G4 of 5 types of VR (variable) genes

**Table 3-B.** Overlap of pathways significantly enriched in GWAS-detected genes and those in the lists T_0_ ∼ T_4_ of 5 types of *exclusive* VR (variable) genes

|  | **gwas.lvmass** | **T<sub>0</sub>** | **T<sub>1</sub>** | **T<sub>2</sub>** | **T<sub>3</sub>** | **T<sub>4</sub>** |
| --- | --- | --- | --- | --- | --- | --- |
| Unique Genes | 1,246 | 243 | 243 | 243 | 243 | 243 |
| Pathways Sig | 2,237 | 111 | 0 | 9 | 1 | 649 |
| Shared Genes |  | 7 | 11 | 16 | 10 | **17** |
| Shared Pathways |  | 9 | 0 | 7 | 0 | **102** |

## DISCUSSION

While hiPSC-derived cell models become increasingly popular for studying human diseases, our knowledge about their fidelity to the original donors’ disease biology is still quite limited. Here we present a novel study to determine sources of transcriptional variabilities in hiPSC-CMs for modeling LVH in the HyperGEN-CiPS study. Our experiments were conducted on hiPSC-CMs obtained from individuals with or without LVH in the HyperGEN cohort, as well as an external donor with a Mendelian form of heart disease. Genome-wide transcriptional variability was assessed with RNA sequencing across hiPSC-CMs, at baseline and following perturbation to the disease-in-a-dish model by stimulation using the potent cardiac drug Endothelin-1 (Carlson et al., 2013). By properly modeling the transcriptional variabilities using generalized linear mixed models, we were able to characterize the sources of observed transcription variations. Careful examination of the variation profiles, both genome-wide and in groups of genes, and linking back to GWAS and eQTL information have led to significant insights about modeling complex human disease using hiPSCs.

<u>First,</u> the variabilities of technical factors are generally smaller than the biological variabilities of interest in hiPSC-CMs. Early studies observed retention of somatic cell memory in hiPSC lines and showed that the problem can be alleviated by prolonged cell culture (later passages) (Bar-Nur et al., 2011; Kim et al., 2010). A more recent study showed that hiPSC lines derived from the same donor are highly similar to each other (Kyttälä et al., 2016). This was verified by another study in the NextGen consortium (Carcamo-Orive et al., 2017) and a meta-analysis combining with data from HipSci Consortium (Germain and Testa, 2017; Kilpinen et al., 2017). A similar observation was reported by (Matsa et al., 2016b) in cardiomyocytes derived from clones across five hiPSC lines. We confirmed and extended that, for a majority of genes, the same is true in our own hiPSC-derived cardiomyocytes. Our analyses showed further that there exist groups of genes for which variabilities due to reprogramming and stimulation may be confounded. This was not surprising because of the genes (and regulators) involved in producing the progeny cardiac cells.

<u>Second,</u> disease state in donors can be manifested as systemic increases of transcriptional variabilities in hiPSC-derived progeny cells. There is a body of literature on the relationship between failures of the transcriptional regulatory systems and diseases in humans (Albert and Kruglyak, 2015; Boyle et al., 2017; Chatterjee et al., 2016; Zhang and Lupski, 2015). Recent studies have taken a systems biology approach and begun to shift focus to the robustness of the transcriptional networks. Network analysis methods were applied to characterize genes and their transcriptional regulators in cardiac dysfunction and other diseases (Nair et al., 2014; Victor et al., 2016). There is a movement to look into the extreme healthy carriers of known disease causal mutations to discover resilient mutations (Friend and Schadt, 2014; Williams, 2016). However, it is quite possible that the resilience or robustness is a feature of the system rather than a single mutation. If that is the case, the hiPSC-derived somatic cell model can provide a useful model for probing the relationship between the transcriptional robustness and resilience to disease.

<u>Third,</u> genes (and more importantly their variable expression) are not created equal in terms of their contribution and presence in an emerging new model system. Factors driving their relevance lie in the controlled experimental conditions or measured disease traits of interest in donors while many others are unobserved (and likely uncontrolled) random or noise factors in the reprogramming process or technical platform employed by the experiment. It is essential that we know which genes might be more sensitive to reprogramming and how their effects may confound with biological effects of interest. It may not be possible for an individual researcher (group) to thoroughly investigate these. However, we should forge a concerted effort in a consortium such as NextGen to establish a venue to cumulatively learn about the nature of such sensitive genes, and catalog them as reference tools to enable improved research science.

<u>Fourth,</u> results from our integrative analyses using eQTL and GWAS analysis results confirmed that the hiPSC-model can be a powerful tool for functional dissection of variants found by large-scale association (genetic epidemiological) studies. One of the original mandates of the NextGen program was to apply the hiPSC technology “to follow up on genomic associations with additional mechanistic information gained from cellular models of disease”. Our results confirm that this is a viable approach. Even though we only used a small portion of our biological samples and did not perform a full-scale eQTL study, indirect usage of publicly available GTEx data helped us discover promising eQTL SNPs from the original GWAS analysis of LVH phenotypes. The GWAS p-values of the 4 functionally annotated SNPs did not reach genome-wide significance level and would be overlooked using a prioritization scheme purely based on statistical test p-values.

Finally, findings from our analyses have additional implications for future hiPSC-derived somatic cells models. For example, the distributional properties of the variable genes in different categories can provide a guideline or reference for power analysis for designing disease-in-a-dish studies. The variable gene lists can provide a starting point to extract markers that may provide means to diagnose problematic issues at particular steps of the reprogramming process of the model. As the field embarks on a new phase of precision medicine, the capability of hiPSC reprogramming technologies to generate donor-specific somatic cells will undoubtedly ignite a new wave of studies employing the disease-in-a-dish models (Biel et al., 2015; Shen et al., 2021), as well as patient oriented drug discovery and drug response/toxicology studies (Davaapil et al., 2020; Li et al., 2017; Neofytou et al., 2015; Sharma et al., 2020). Combining the power of this new technology with the rich information packed in the numerous existing (and oncoming) GWAS and other types of genomic data, additional research is sorely needed for better understanding the fidelity of new models both qualitatively and quantitatively. Our findings about transcriptional variabilities and landscape in hiPSC-CMs have confirmed the validity of this model, while raising some new questions (e.g., about the variable gene lists and how to separate confounding effects). Answering these questions will demand further research on how to improve our collective understanding of the problem, and will have important implication for future hiPSC-based disease-in-a-dish models.

## AUTHOR CONTRIBUTIONS

Conceptualization and design of study, U.B, C.C.G; hiPSC production, FCDI; hiPSC-CM generation, RNA-seq experiments, A.M, A.T; Data curation, modeling and analysis, P.A., C.C.G, W.Y., X.S.; Manuscript writing: C.C.G; Review and editing: A.T, B.A, P.A., A.M., D.A.K., U.B. Funding Acquisition: U.B.

## ACKNOWLEDGEMENTS

This research was supported in part by NIH grants HL107437, HL125580, and HL055673.

## EXPERIMENTAL PROCEDURES

### Generation of hiPSC-CMs & other experimental data

This “clone experiment” was conducted using a subsample from the ongoing HyperGEN-CiPS Study (Functional GWAS for LVH Using hiPSC-derived Cardiomyocytes). The HyperGEN-CiPS is part of the NextGen Program funded by the NHLBI (National Heart, Lung, and Blood Institute) aimed at creating hiPSC libraries and progeny somatic cell lines to advance functional dissection of the genetic architecture underlying diseases relevant to NHLBI’s missions (Warren et al., 2017). It is also a sub-study of HyperGEN (Hypertension Genetic Epidemiology Network), a family-based cohort for studying genetic causes of hypertension and related diseases in European and African Americans (EA & AA). The original HyperGEN study recruited hypertensive sibships, along with their normotensive adult offspring and an age-matched random sample. In the HyperGEN-CiPS study, blood samples and other phenotype measurements were collected in a subsample of 250 selected from the original HyperGEN cohorts to study the molecular changes associated with the development of left ventricular hypertrophy (LVH) by analyzing expression changes under various conditions in hiPSC-CM cell lines. One goal is to functionally dissect and annotate disease variants found by genome-wide association analysis (GWAS). The cell lines were generated from donors in the HyperGEN-CiPS study, with normal or high left ventricular mass (LVmass) (Lang et al., 2015). Batches of hiPSC-CMs were generated from hiPSC clones derived from the same and different donors. A cardiac stimulant (ET-1, i.e. endothelin-1) was applied to the hiPSC-CMs to trigger hypertrophic responses and RNA-seq was used to measure transcriptional changes. Using a PIGGY-BACK reprogramming approach, we generated hiPSCs from 248 individuals with genome-wide genotype data and exquisite phenotypic data, and subsequently have derived cardiomyocytes. For the analyses reported *here*, we have conducted 50 experiments in 25 clones (21 unique ones) derived from 18 hiPSC cell lines made in 6 individual HyperGEN donors and one case of external sample of familial hypertrophic cardiomyopathy. Additional Information can be found online, including extensive descriptions of the study samples and all experimental and statistical analysis methods and procedures, in the Supplemental Experimental Procedures section.

## Data Availability

https://zenodo.org/records/10001344

## REFERENCES

Albert, F.W., and Kruglyak, L. (2015). The role of regulatory variation in complex traits and disease. Nat Rev Genet 16, 197–212.

Bar-Nur, O., Russ, H.A., Efrat, S., and Benvenisty, N. (2011). Epigenetic memory and preferential lineage-specific differentiation in induced pluripotent stem cells derived from human pancreatic islet beta cells. Cell Stem Cell 9, 17–23.

Biel, N.M., Santostefano, K.E., DiVita, B.B., El Rouby, N., Carrasquilla, S.D., Simmons, C., Nakanishi, M., Cooper-DeHoff, R.M., Johnson, J.A., and Terada, N. (2015). Vascular Smooth Muscle Cells From Hypertensive Patient-Derived Induced Pluripotent Stem Cells to Advance Hypertension Pharmacogenomics. Stem Cells Transl Med 4, 1380–1390.

Boyle, E.A., Li, Y.I., and Pritchard, J.K. (2017). An Expanded View of Complex Traits: From Polygenic to Omnigenic. Cell 169, 1177–1186.

Braam, S.R., Tertoolen, L., van de Stolpe, A., Meyer, T., Passier, R., and Mummery, C.L. (2010). Prediction of drug-induced cardiotoxicity using human embryonic stem cell-derived cardiomyocytes. Stem Cell Res 4, 107–116.

Carcamo-Orive, I., Hoffman, G.E., Cundiff, P., Beckmann, N.D., D’Souza, S.L., Knowles, J.W., Patel, A., Papatsenko, D., Abbasi, F., Reaven, G.M., et al. (2017). Analysis of Transcriptional Variability in a Large Human iPSC Library Reveals Genetic and Non-genetic Determinants of Heterogeneity. Cell Stem Cell 20, 518–532 e519.

Carlson, C., Koonce, C., Aoyama, N., Einhorn, S., Fiene, S., Thompson, A., Swanson, B., Anson, B., and Kattman, S. (2013). Phenotypic screening with human iPS cell-derived cardiomyocytes: HTS-compatible assays for interrogating cardiac hypertrophy. J Biomol Screen 18, 1203–1211.

Chatterjee, S., Kapoor, A., Akiyama, J.A., Auer, D.R., Lee, D., Gabriel, S., Berrios, C., Pennacchio, L.A., and Chakravarti, A. (2016). Enhancer Variants Synergistically Drive Dysfunction of a Gene Regulatory Network In Hirschsprung Disease. Cell 167, 355–368 e310.

Davaapil, H., Shetty, D.K., and Sinha, S. (2020). Aortic "Disease-in-a-Dish": Mechanistic Insights and Drug Development Using iPSC-Based Disease Modeling. Front Cell Dev Biol 8, 550504.

Dell’Era, P., Benzoni, P., Crescini, E., Valle, M., Xia, E., Consiglio, A., and Memo, M. (2015). Cardiac disease modeling using induced pluripotent stem cell-derived human cardiomyocytes. World J Stem Cells 7, 329–342.

Friend, S.H., and Schadt, E.E. (2014). Translational genomics. Clues from the resilient. Science 344, 970–972.

Germain, P.L., and Testa, G. (2017). Taming Human Genetic Variability: Transcriptomic Meta-Analysis Guides the Experimental Design and Interpretation of iPSC-Based Disease Modeling. Stem Cell Reports 8, 1784–1796.

Guenther, M.G., Frampton, G.M., Soldner, F., Hockemeyer, D., Mitalipova, M., Jaenisch, R., and Young, R.A. (2010). Chromatin structure and gene expression programs of human embryonic and induced pluripotent stem cells. Cell Stem Cell 7, 249–257.

Kehat, I., Kenyagin-Karsenti, D., Snir, M., Segev, H., Amit, M., Gepstein, A., Livne, E., Binah, O., Itskovitz-Eldor, J., and Gepstein, L. (2001). Human embryonic stem cells can differentiate into myocytes with structural and functional properties of cardiomyocytes. J Clin Invest 108, 407–414.

Kilpinen, H., Goncalves, A., Leha, A., Afzal, V., Alasoo, K., Ashford, S., Bala, S., Bensaddek, D., Casale, F.P., Culley, O.J., et al. (2017). Common genetic variation drives molecular heterogeneity in human iPSCs. Nature 546, 370–375.

Kim, K., Doi, A., Wen, B., Ng, K., Zhao, R., Cahan, P., Kim, J., Aryee, M.J., Ji, H., Ehrlich, L.I.R., et al. (2010). Epigenetic memory in induced pluripotent stem cells. Nature 467, 285–290.

Knollmann, B.C. (2013). Induced pluripotent stem cell-derived cardiomyocytes: boutique science or valuable arrhythmia model? Circ Res 112, 969–976; discussion 976.

Kyttälä, A., Moraghebi, R., Valensisi, C., Kettunen, J., Andrus, C., Pasumarthy, K.K., Nakanishi, M., Nishimura, K., Ohtaka, M., Weltner, J., et al. (2016). Genetic Variability Overrides the Impact of Parental Cell Type and Determines iPSC Differentiation Potential. Stem Cell Reports 6, 200–212.

Lang, R.M., Badano, L.P., Mor-Avi, V., Afilalo, J., Armstrong, A., Ernande, L., Flachskampf, F.A., Foster, E., Goldstein, S.A., Kuznetsova, T., et al. (2015). Recommendations for cardiac chamber quantification by echocardiography in adults: an update from the American Society of Echocardiography and the European Association of Cardiovascular Imaging. J Am Soc Echocardiogr 28, 1–39 e14.

Law, C.W., Chen, Y., Shi, W., and Smyth, G.K. (2014). voom: Precision weights unlock linear model analysis tools for RNA-seq read counts. Genome Biol 15, R29.

Li, Y., Sallam, K., Schwartz, P.J., and Wu, J.C. (2017). Patient-Specific Induced Pluripotent Stem Cell-Based Disease Model for Pathogenesis Studies and Clinical Pharmacotherapy. Circ Arrhythm Electrophysiol 10.

Matsa, E., Ahrens, J.H., and Wu, J.C. (2016a). Human induced pluripotent stem cells as a platform for personalized and precision cardiovascular medicine. Physiological Reviews 96, 1093–1126.

Matsa, E., Burridge, P.W., Yu, K.H., Ahrens, J.H., Termglinchan, V., Wu, H., Liu, C., Shukla, P., Sayed, N., Churko, J.M., et al. (2016b). Transcriptome Profiling of Patient-Specific Human iPSC-Cardiomyocytes Predicts Individual Drug Safety and Efficacy Responses In Vitro. Cell Stem Cell 19, 311–325.

McIntyre, L.M., Lopiano, K.K., Morse, A.M., Amin, V., Oberg, A.L., Young, L.J., and Nuzhdin, S.V. (2011). RNA-seq: technical variability and sampling. BMC Genomics 12, 293.

Molecular Signatures Database v6.0. http://software.broadinstitute.org/gsea/msigdb, Accessed: 2017.

Nair, J., Ghatge, M., Kakkar, V.V., and Shanker, J. (2014). Network analysis of inflammatory genes and their transcriptional regulators in coronary artery disease. PLoS One 9, e94328.

Neofytou, E., O’Brien, C.G., Couture, L.A., and Wu, J.C. (2015). Hurdles to clinical translation of human induced pluripotent stem cells. J Clin Invest 125, 2551–2557.

Onder, T.T., and Daley, G.Q. (2012). New lessons learned from disease modeling with induced pluripotent stem cells. Curr Opin Genet Dev 22, 500–508.

Risso, D., Ngai, J., Speed, T.P., and Dudoit, S. (2014). Normalization of RNA-seq data using factor analysis of control genes or samples. Nat Biotechnol 32, 896–902.

Ritchie, M.E., Phipson, B., Wu, D., Hu, Y., Law, C.W., Shi, W., and Smyth, G.K. (2015). Limma powers differential expression analyses for RNA-sequencing and microarray studies. Nucleic Acids Research 43, e47.

Rouhani, F., Kumasaka, N., de Brito, M.C., Bradley, A., Vallier, L., and Gaffney, D. (2014). Genetic Background Drives Transcriptional Variation in Human Induced Pluripotent Stem Cells. PLoS Genetics 10.

Sharma, A., Sances, S., Workman, M.J., and Svendsen, C.N. (2020). Multi-lineage Human iPSC-Derived Platforms for Disease Modeling and Drug Discovery. Cell Stem Cell 26, 309–329.

Shen, M., Quertermous, T., Fischbein, M.P., and Wu, J.C. (2021). Generation of Vascular Smooth Muscle Cells From Induced Pluripotent Stem Cells: Methods, Applications, and Considerations. Circ Res 128, 670–686.

Sinnecker, D., Laugwitz, K.L., and Moretti, A. (2014). Induced pluripotent stem cell-derived cardiomyocytes for drug development and toxicity testing. Pharmacol Ther 143, 246–252.

Smyth, G.K. (2005). Limma: linear models for microarray data. In Bioinformatics and Computational Biology Solutions using R and Bioconductor, R. Gentleman, W. Huber, V. Carey, and R.A. Irizarry, eds. (New York: Springer-Verlag).

Subramanian, A., Tamayo, P., Mootha, V.K., Mukherjee, S., Ebert, B.L., Gillette, M.A., Paulovich, A., Pomeroy, S.L., Golub, T.R., Lander, E.S., et al. (2005). Gene set enrichment analysis: A knowledge-based approach for interpreting genome-wide expression profiles. Proceedings of the National Academy of Sciences of the United States of America 102, 15545–15550.

Takahashi, K., and Yamanaka, S. (2006). Induction of pluripotent stem cells from mouse embryonic and adult fibroblast cultures by defined factors. Cell 126, 663–676.

Tanaka, A., Yuasa, S., Node, K., and Fukuda, K. (2015). Cardiovascular Disease Modeling Using Patient-Specific Induced Pluripotent Stem Cells. Int J Mol Sci 16, 18894–18922.

Tapia, N., and Schöler, H.R. (2016). Molecular Obstacles to Clinical Translation of iPSCs. Cell Stem Cell 19, 298–309.

Vasan, R.S., Larson, M.G., Aragam, J., Wang, T.J., Mitchell, G.F., Kathiresan, S., Newton-Cheh, C., Vita, J.A., Keyes, M.J., O’Donnell, C.J., et al. (2007). Genome-wide association of echocardiographic dimensions, brachial artery endothelial function and treadmill exercise responses in the Framingham Heart Study. BMC Med Genet 8 *Suppl 1*, S2.

Victor, J.M., Debret, G., Lesne, A., Pascoe, L., Carrivain, P., Wainrib, G., and Hugot, J.P. (2016). Network Modeling of Crohn’s Disease Incidence. PLoS One 11, e0156138.

Warren, C.R., O’Sullivan, J.F., Friesen, M., Becker, C.E., Zhang, X., Liu, P., Wakabayashi, Y., Morningstar, J.E., Shi, X., Choi, J., et al. (2017). Induced Pluripotent Stem Cell Differentiation Enables Functional Validation of GWAS Variants in Metabolic Disease. Cell Stem Cell 20, 547–557 e547.

Williams, S.C. (2016). Genetic mutations you want. Proc Natl Acad Sci U S A 113, 2554–2557.

Yu, J., Vodyanik, M.A., Smuga-Otto, K., Antosiewicz-Bourget, J., Frane, J.L., Tian, S., Nie, J., Jonsdottir, G.A., Ruotti, V., Stewart, R., et al. (2007). Induced pluripotent stem cell lines derived from human somatic cells. Science 318, 1917–1920.

Zhang, F., and Lupski, J.R. (2015). Non-coding genetic variants in human disease. Hum Mol Genet 24, R102–110.

