## Supplemental text, tables and figures for "Transcriptional Variabilities in Human hiPSC-derived Cardiomyocytes: All Genes Are Not Equal and Their Robustness May Foretell Donor’s Disease Susceptibility"

[1]Institute for Informatics, Data Science and Biostatistics, [2]Department of Genetics, Washington University School of Medicine, St. Louis, MO; [3]Department of Pediatrics, Medical College of Wisconsin, Milwaukee, WI; [4]Department of Genetic Medicine, Weill Cornell Medicine, Doha, Qatar and the Division of Epidemiology, University of Utah School of Medicine, Salt Lake City, UT; [5]Division of Preventive Medicine, Department of Medicine, University of Alabama at Birmingham, AL; [6]Office of the Provost, University of South Carolina, Columbia, SC; [7]FujiFilm Cellular Dynamics International (FCDI), Madison WI; [8]University of Washington, Department of Pathology, Seattle, WA

#### SUPPLEMENTAL EXPERIMENTAL PROCEDURES

**Study donor samples:** This "clone experiment" was conducted using a subsample from the ongoing HyperGEN-CiPS Study (Functional GWAS for LVH Using hiPSC-derived Cardiomyocytes). The HyperGEN-CiPS is part of the NextGen Program funded by the NHLBI (National Heart, Lung, and Blood Institute) aimed at creating hiPSC libraries and progeny somatic cell lines to advance functional dissection of the genetic architecture underlying diseases relevant to NHLBI's missions (Warren et al., 2017). It is also a sub-study of HyperGEN (Hypertension Genetic Epidemiology Network), a family-based cohort for studying genetic causes of hypertension and related diseases in European and African Americans (EA & AA). The original HyperGEN study is a multicenter study in which hypertensive sibships diagnosed prior to age 60 were enrolled from four separate field centers from 1995 to 2005 (Williams et al., 2000). Hypertension was defined as current antihypertensive medication use or having an average systolic blood pressure ≥140 mm Hg and/or diastolic blood pressure ≥90 mm Hg measured at two separate clinic visits. Individuals with a history of type 1 diabetes or severe renal disease were excluded. In the HyperGEN-CiPS study, blood samples and other phenotype measurements were collected or re-measured at the UAB & UTH field centers in a subsample of 250 selected from the original HyperGEN cohorts. In order to validate the identity of the hiPSC donors, the iPSC clones were genotyped using Illumina HumanCore24 and the results were compared to their respective donors' germline genotypes in the original GWAS dataset in HyperGEN. Because different genotyping platforms (Afftmetrix 5.0 and Affymetrix 6.0 arrays) were used by HyperGEN GWAS, we first identified the subset of genotyping variants/probes measured by both platforms with the same reference alleles. This gave us a total of 18,756 probes, over which the genotype calls between the hiPSC clones and their donors were compared using an in-house pipeline scripted in R and Python.

**Donor data:** *HyperGEN phenotyping:* Baseline phenotype and covariates data were collected by the original HyperGEN as previously described. These include echocardiographic measures collected using Doppler, two-dimensional (2D), and M-mode (2D-guided) echocardiograms following a standardized protocol previously described (Devereux and Roman, 1995). M-mode and 2D echocardiograms via the parasternal acoustic window were recorded for ≥10 beats. Measurements were made at the HyperGEN echocardiography reading center using a computerized review station equipped with a digitizing tablet and monitor overlay used for calibration and quantification (Digisonics, Inc., Houston, Texas). LV linear dimensions were measured by M-mode or 2D echocardiography according to American Society of Echocardiography recommendations (Sahn et al., 1978; Schiller et al., 1989). LV mass was calculated using end-diastolic dimensions by an anatomically validated formula (Devereux et al., 1984), and end-diastolic and end-systolic LV volumes calculated by the Teichholz method (Teichholz et al., 1976) were used to calculate LV ejection fraction (Kizer et al., 2004).

*HyperGEN genotyping & GWAS findings:* The original HyperGEN study has QC'ed GWAS data genotyped using the Affymetrix 6.0 and/or 5.0 arrays, in 1,258 (1,083+175, 6.0+5.0) African American and 1,270 (all 5.0) Caucasian American Subjects,. After extensive QC (missing rate, MAF, Hardy-Weinberg, Mendelian errors), the cohorts retained 837,134 (in AA) and 358,327 autosomal SNPs (in CA), respectively, for conventional GWAS analyses. Findings from GWAS analyses of LVH related echo traits were reported elsewhere (Arnett et al., 2011). We only used limited results for annotation/discussion of our findings in the present study. In the present report, we used the GWAS results found for 2 LVH traits, logarithm of LV mass (LVM) and ejection fraction (EF), using Merlin (Abecasis et al., 2002) with score test option (--fastAssoc) and packages in statistical software **R**.

**Generation of Individual hiPSC-CM Cell Lines**

The generation of the parent iPSCs and differentiation to Cardiomyocytes (FujiFilm Cellular Dynamics International, FCDI) were done as described previously (Ma et al., 2011). Reprogramming of peripheral blood monocytes (PBMCs) for 124 African-American donor samples from the HyperGEN cohort was performed with modification of the episomal method described by Yu and colleagues (Yu et al., 2011). In brief, PBMCs were collected from standard blood draws and expanded for approximately eight days in Iscove's Modified Dulbecco's Medium (IMDM; ThermoFisher Scientific) with supplements. Expanded PBMCs were then transfected with the OCT4, SOX2, NANOG, LIN28, L-MYC, KLF4 and SV40LT transgenes as described previously (Yu et al., 2009). The iPSC chromosomal integrity was checked with SNP array (Infinium Human Core Bead Chip; Illumina). Donor specific iPSCs were then prepared for subsequent cardiomyocyte differentiation and selection procedures similar to iCell Cardiomyocytes (Ma et al., 2011). The iPSC clones were engineered to exhibit cardiomyocyte-specific blasticidin resistance by inserting the coding region of the blasticidin S deaminase (*BSD*, *Aspergillus terreus)* downstream of the last exon of the native *MYH6* gene coding region through nuclease-mediated homologous recombination. A picornavirus 2A translational slip site was inserted between the BSD and *MYH6* coding regions to prevent haplo-insufficiency of the myosin heavy chain (de Felipe et al., 2003; Donnelly et al., 2001). Resulting iPSC colonies were subjected to genomic DNA PCR screening to correctly identify targeted homologous recombinants using primer sets that spanned both arms of the recombination site. Cardiomyocytes at greater than 90% purity were derived through the differentiation and selection procedures described by Ma and colleagues (Ma et al., 2011). Cardiomyocytes were matched to the donor material through short-tandem repeat (STR) ID testing.

The 21 unique hiPSC-CMs were plated at $2.0 \times 10^4$ cells/well in 96-well plates pre-coated with fibronectin (day 0) and cultured for 14 days post plating. At day 2, the cells were switched from plating media to iCell Cardiomyocyte Maintenance Medium (Cellular Dynamics International). After an additional 7 days of recovery (day 9), cells were then switched to William's E medium supplemented with Cocktail B (1:25) from the Hepatocyte Maintenance Supplement Pack (SWE) (ThermoFisher Scientific). At day 13 post plating, cells were either stimulated for 18 hours with ET-1 (Sigma Aldrich) at $10^{-8}$M or received no ET-1 stimulation.

Eighteen hours post ET-1 stimulation cells were harvested with Total RNA Purification 96-Well Kit (Norgen Biotek Corp.). Total RNA was extracted per manufacturer's recommendations, resuspended in nuclease-free water. The RNA used for library preparation was then further concentrated using Qiagen RNeasy MinElute Cleanup Kit (Qiagen) and quantified by UV spectrophotometry (NanoDrop™ 2000, Thermo Scientific).).

**RNA Sequencing:**

Barcoded Whole Transcriptome RNA libraries from 1.5ug of rRNA depleted total RNA (Ribo-Zero rRNA removal Kit, Epicentre) were prepared using Ion Torrent Total RNA Seq-Kit for the ABI Library Builder™ System as per manufacturer's instructions (Thermo Fisher Scientific). Libraries were assessed for quality and quantified using Agilent Bioanalyzer High sensitivity chip (Agilent Technologies, Inc). Libraries were then prepared and sequenced using the Ion PI™ Template OT2 200 Kit v3 on the Ion Torrent OneTouch2 system (Thermo Fisher Scientific) following manufactures instruction. Single-end sequencing was performed using the Ion Torrent Proton sequencing platform following manufacturer's instructions.

**RNA-Seq Pre-processing:**

RNA-Seq reads were aligned to human reference genome build hg19 using RNA-STAR. Reads overlapping gene regions were counted using htseq-count function from the HTSeq Python package. For RNA-seq data pre-processing and quality assessment, counts per million (cpm) and TMM normalization (Robinson and Oshlack, 2010) implemented in edgeR (Robinson et al., 2010) were used. Transcripts (genes) with cpm$\geq$1 in at least one experiment ore more were retained for further analysis. For QA analyses, we applied TMM normalization (Robinson and Oshlack, 2010) implemented in edgeR (Robinson et al., 2010) for normalization and GLM to correct for the effects of stimulation experiment. The procedure RUVg implemented in the RUVseq package (Risso et al., 2014) was applied to remove hidden "unwanted" variations using empirical negative control genes that were least variable as identified by the GLM. It was then followed by visual examination of Box plots of expression levels across all 50 experiments and plots of the first 2 principal components extracted from the so-normalized expression matrix.

To check sample clustering, we applied voom (Law et al., 2014) implemented in limma (Ritchie et al., 2015; Smyth, 2005) (Smyth, 2005) and included experiment batch and other covariates that might be indicators of experiment quality including the percentage of reads successfully mapped, and the number of cells that survive the plating process (post plate counts). Expression residuals from the voom normalization were used for clustering analysis to assess quality of the experiments for further statistical analysis.

**Statistical Analysis:**

**Characterize sources of variations in expression levels**:

*Statistical Models:* To characterize the different sources of expression variability, we need to properly model the random factors using the replicate samples. Following the argument used by edgeR (McCarthy et al., 2012; Robinson et al., 2010) that total expression variation can be decomposed into technical and biological variations following Poisson and gamma distributions respectively, we model the observed expression gene counts data using a negative binomial distribution. To accommodate the fact that the design of the clone experiment was unbalanced, we used generalized linear mixed models (GLMM) implemented in the R package lme4. Five random effects were included to capture variations due to: RNA-seq, CMs clones generation, hiPSC lines derivation, donor/host individual biology, and ET-1 stimulation experiment. We considered several models by incrementally introducing these random effects (see Figure 1B), where the first 4 random effects were nested in a sequential manner. Three technical factors were always included as fixed effects: RNA-seq batch, post plate counts, and percentage of reads successfully mapped. The models were implemented in R using `lme4::glemr.nb()` to fit GLMM for the negative binomial family.

*Evaluate contributions of technical and biological effects:* Variance components estimated by the fitted GLMM were extracted for all converged models. Proportions of the variances relative to the combined variation due to all 5 random effects were estimated to study contributions of the various technical and biological effects. The violin plot (Hintze and Nelson, 1998) combining a boxplot and a rotated kernel density plot on each side is used to visually examine the summary characteristics of data (median, inner quartile range [IQR], etc.) conveyed by the boxplot together with the concentrations of density as indicated by the widths on the "violin". R package *`vioplot`* was used to make the violin plots. The estimated variances were used to compare "signal" variabilities due to biological effects of ET-1 stimulation (*group*) and donor biology (*id*), to "noise" ones due to (undesirable) technical noises of RNA-seq (*tech*), CMs generation (*cms*) and hiPSC derivation (*hiPSC*). For this purpose, variance components were examined both individually and collectively for an exhaustive list of combinations.

**Study the "genetic robustness" hypothesis of complex disease**:

For complex disease such as LVH, the "genetic robustness" hypothesis proposes that, in one way or another, relevant components of biological systems in disease patients are compromised and render the systems less robust under stress. Consequently, perturbations by drug stimulation would result in higher *variability* of gene expression in hiPSC-CMs obtained from LVH than from normal donors. We verify this in 2 complimentary manners. First, we compare variances of expression in hiPSC-CMs derived from LVH donors to those from normal subjects, where a ratio $>1$ indicates that the gene is more sensible to perturbation in a system compromised by LVH. The genome-wide distribution of the ratios will then provide corroborative evidence for the robustness hypothesis if increased variability is observed in many genes and/or in groups of genes that are relevant to heart health. Second, we compare the numbers of genes significantly differentially expressed (DE genes) between LVH- vs normal-donor hiPSC-CMs in stimulated cell lines with those in unstimulated ones, and the numbers of DE genes between stimulated vs baseline in LVH-donor with those in normal-donor cells. To enable the comparisons, we re-fitted nested linear models on the GLMM normalized expression values using 2 nested models: with the donor's LVH status nested within stimulation condition, and *vice versa*. If the robustness hypothesis is valid then we should observe both more *LVH-vs-normal* DE genes in stimulated cells, and more *stimulated-vs-baseline* DE genes in LVH-donor hiPSC-CMs clones.

**Variable (VR) genes sensitive to different steps of a disease-in-a-dish experiment:**

Empirical data shows that genes are not made equal when come to reacting to factors involved in a hiPSC-based experiment. It is important to distinguish those genes that are more prone to noise factors in such experiments, and to correctly characterize at which step(s) such a sensitive gene becomes *undesirably* more variable. Using estimates from the GLMM modeling, we evaluated 3 ways to characterize such VR gene groups by: (1) likelihood ration test (LRT) contrasting models included in Figure 1B; (2) one-side thresholds on quantiles of estimated variances of the random effects; (3) two-side thresholds on the quantiles. More specifically, for (1) we performed a LRT with 3 degrees of freedom comparing model #65 to #62 for genes variable because of "noises" in RNA sequencing, CM cloning, or hiPSC generation (*"noise" VR genes*); a LRT with 2 degrees of freedom comparing model #62 to #60 for genes variable because of "signals" either via stimulation or individual biology, both connected to donors' LVH status (*"signal" VR gene*s); and a series of LRTs of 1 degree of freedom comparing adjacent models in Figure 1B for VR genes due to a single factor. For (2) we put a threshold on the quantiles of their variance components estimated by the restricted model #65. Using the number of 62 noise genes obtained by LRT that turned out being mostly attributable to hiPSC generation, we projected that there would be about 310 (=62*5) gene in each type of VR genes, which translated to a quantile threshold of top 1.8%. We then generate 5 types of VR genes, each due to a random effect in model #65, by applying the quantile threshold to the relevant variance components (i.e., Var(*type*) ≥ 98.2-percentile, for the "*type*" random effect) to obtain ~310 genes in each list. For (3) we apply 2 thresholds on the quantiles of variances: one on the top X-percentile with respect to a given type of variability, the other on the bottom Y-percentile of the sum of variances due to the remaining 4 other random effects. This ensures the resulting VR genes are sensitive only to the single given random effect.

*Evaluate source-specific variabilities by violin plots:* For each random effect, we calculated the fraction of variance due to the factor, and created violin-plots (Hintze and Nelson, 1998) as described above. Each plot demonstrates where the contribution to expression variabilities is concentrated for the corresponding random factor. If the VR genes selected do a good job in capturing variabilities due to the intended random factor, then one should see the plot with an elevated non-zero median for that factor while the plots for other factors have median of (or very close to) zero and little density elsewhere.

**Cross validation of genes responsive to ET-1 stimulation:**

*Cross validation by GWAS hits & eQTLs:* A type of validation of the list of VR genes responsive to ET-1 stimulation is provided by co-localization of LVH GWAS-detected SNPs and expression QTLs in LVH relevant tissues that regulates any of the genes in the list. The eQTL data from the Genotype-Tissue Expression (GTEx) Project (V.6) and the results from HyperGEN GWAS analyses of the left ventricular (LV) mass and the ejection fraction (EF) were used for the co-localization analysis. For any given gene, we extracted eQTL SNPs from GTEx detected at the significance level of $\alpha=1.0E\text{-}5$ in any of 4 LVH related tissues (left ventricle, atrial appendage, coronary artery, and aorta) analyzed by GTEx. Significant GWAS-found SNPs at a threshold of $\alpha=1.0E\text{-}3$ were extracted from previous HyperGEN GWAS analyses [(Arnett et al., 2011) and unpublished data] for 2 LVH traits of left ventricular (LV) mass and ejection fraction (EF). Latest annotation provided by the manufacture (GenomeWideSNP_6.na35.annot) for SNP locations and associated genes were used to construct a map of the GWAS hits. Co-localization was then analyzed by merging the GTEx derived eQTL SNPs for the ET-1 responsive genes with the map of the HyperGEN GWAS hits for the 2 LVH traits. Finally, effect sizes of co-localized eQTL SNPs in (up to) 43 tissues as estimated by GTEx were extracted and plotted to visually examine for their potential in LVH related tissues.

*Enrichment analysis of known pathways in lists of VR genes:* Another way to assess the relevance of the VR genes responsive to ET-1 stimulation is to examine sharing of common pathways or gene sets that are significantly enriched in the VR gene list and among genes harboring the GWAS hits. If the VR genes for stimulation are indeed important to LVH, then we should see many enriched pathways in the group also significantly enriched among GWAS-hit genes. Routines in the Python package SciPy were used to perform hypergeometric test of each of the 18,026 pathways/gene sets in the Molecular Signatures Database (MSigDB; Subramanian et al., 2005) for significant enrichment of the gene set in any given list of VR genes. P-values were adjusted for multiple comparisons using the Benjamini-Hochberg method (Benjamini and Hochberg, 1995) to control false discovery rates (FDR). Significantly enriched pathways/gene sets were obtained using FDR cutoff of 0.05. Also reported were the numbers of shared genes and common significantly enriched pathways between GWAS-hit genes and the target VR gene list. The same method can be applied to verify sharing of significantly enriched pathways between the GWAS-hit genes and other types of VR gene lists. Comparing the results across the VR gene lists can provide further evidence for (or against) the relevance of the VR genes responsive to ET-1 stimulation.

**Computer software and packages:**

Except when stated otherwise, all analyses were performed using R version 3.3.0 and packages (see Resources Table in Supplementary Method). Mapping of SNPs to reference genomes used either manufacturer's annotation (GWAS array) or NCBI databases via R packages `biomaRt` and `reutils`. RNA-seq data analyses used R packages `edgeR`, `limma` and `DESeq2`. Hierarchical clustering was done using `hclust` and plotted by `dendextend` & `circlize`. Heatmaps were drawn by `pheatmap`.

#### SUPPLEMENTAL REFERENCES

Abecasis, G.R., Cherny, S.S., Cookson, W.O., and Cardon, L.R. (2002). Merlin--rapid analysis of dense genetic maps using sparse gene flow trees. Nat Genet *30*, 97-101.

Arnett, D.K., Meyers, K.J., Devereux, R.B., Tiwari, H.K., Gu, C.C., Vaughan, L.K., Perry, R.T., Patki, A., Claas, S.A., Sun, Y.V.*, et al.* (2011). Genetic variation in NCAM1 contributes to left ventricular wall thickness in hypertensive families. Circ Res *108*, 279-283.

Benjamini, Y., and Hochberg, Y. (1995). Controlling the False Discovery Rate: a practical and powerful approach to multiple testing. Journal of the Royal Statistical Society, Series B *57*, 289-300.

de Felipe, P., Hughes, L.E., Ryan, M.D., and Brown, J.D. (2003). Co-translational, intraribosomal cleavage of polypeptides by the foot-and-mouth disease virus 2A peptide. J Biol Chem *278*, 11441-11448.

Devereux, R.B., Lutas, E.M., Casale, P.N., Kligfield, P., Eisenberg, R.R., Hammond, I.W., Miller, D.H., Reis, G., Alderman, M.H., and Laragh, J.H. (1984). Standardization of M-mode echocardiographic left ventricular anatomic measurements. J Am Coll Cardiol *4*, 1222-1230.

Devereux, R.B., and Roman, M.J. (1995). Evaluation of cardiac function and vascular structure and function by echocardiography and other noninvasive techniques. In Hypertension: pathophysiology, diagnosis, and management, J.H. Laragh, and B.M. Brenner, eds. (New York: Raven P), pp. 1969-1985.

Donnelly, M.L., Luke, G., Mehrotra, A., Li, X., Hughes, L.E., Gani, D., and Ryan, M.D. (2001). Analysis of the aphthovirus 2A/2B polyprotein 'cleavage' mechanism indicates not a proteolytic reaction, but a novel translational effect: a putative ribosomal 'skip'. The Journal of general virology *82*, 1013-1025.

Hintze, J.L., and Nelson, R.D. (1998). Violin plots: a box plot-density trace synergism. The American Statistician *52*, 181-184.

Kizer, J.R., Arnett, D.K., Bella, J.N., Paranicas, M., Rao, D.C., Province, M.A., Oberman, A., Kitzman, D.W., Hopkins, P.N., Liu, J.E.*, et al.* (2004). Differences in left ventricular structure between black and white hypertensive adults: the Hypertension Genetic Epidemiology Network study. Hypertension *43*, 1182-1188.

Law, C.W., Chen, Y., Shi, W., and Smyth, G.K. (2014). voom: Precision weights unlock linear model analysis tools for RNA-seq read counts. Genome Biol *15*, R29.

Ma, J., Guo, L., Fiene, S.J., Anson, B.D., Thomson, J.A., Kamp, T.J., Kolaja, K.L., Swanson, B.J., and January, C.T. (2011). High purity human-induced pluripotent stem cell-derived cardiomyocytes: electrophysiological properties of action potentials and ionic currents. American Journal of Physiology Heart and Circulatory Physiology *301*, H2006-2017.

McCarthy, D.J., Chen, Y., and Smyth, G.K. (2012). Differential expression analysis of multifactor RNA-Seq experiments with respect to biological variation. Nucleic Acids Res *40*, 4288-4297.

Molecular Signatures Database v6.0. http://software.broadinstitute.org/gsea/msigdb, Accessed: 2017.

Risso, D., Ngai, J., Speed, T.P., and Dudoit, S. (2014). Normalization of RNA-seq data using factor analysis of control genes or samples. Nat Biotechnol *32*, 896-902.

Ritchie, M.E., Phipson, B., Wu, D., Hu, Y., Law, C.W., Shi, W., and Smyth, G.K. (2015). Limma powers differential expression analyses for RNA-sequencing and microarray studies. Nucleic Acids Research *43*, e47.

Robinson, M.D., McCarthy, D.J., and Smyth, G.K. (2010). edgeR: A Bioconductor package for differential expression analysis of digital gene expression data. Bioinformatics *26*, 139-140.

Robinson, M.D., and Oshlack, A. (2010). A scaling normalization method for differential expression analysis of RNA-seq data. Genome Biology *11*.

Sahn, D.J., DeMaria, A., Kisslo, J., and Weyman, A. (1978). Recommendations regarding quantitation in M-mode echocardiography: results of a survey of echocardiographic measurements. Circulation *58*, 1072-1083.

Schiller, N.B., Shah, P.M., Crawford, M., DeMaria, A., Devereux, R., Feigenbaum, H., Gutgesell, H., Reichek, N., Sahn, D., Schnittger, I., *et al.* (1989). Recommendations for quantitation of the left ventricle by two-dimensional echocardiography. American Society of Echocardiography Committee on Standards, Subcommittee on Quantitation of Two-Dimensional Echocardiograms. J Am Soc Echocardiogr *2*, 358-367.

Smyth, G.K. (2005). Limma: linear models for microarray data. In Bioinformatics and Computational Biology Solutions using R and Bioconductor, R. Gentleman, W. Huber, V. Carey, and R.A. Irizarry, eds. (New York: Springer-Verlag).

Subramanian, A., Tamayo, P., Mootha, V.K., Mukherjee, S., Ebert, B.L., Gillette, M.A., Paulovich, A., Pomeroy, S.L., Golub, T.R., Lander, E.S., *et al.* (2005). Gene set enrichment analysis: A knowledge-based approach for interpreting genome-wide expression profiles. Proceedings of the National Academy of Sciences of the United States of America *102*, 15545-15550.

Teichholz, L.E., Kreulen, T., Herman, M.V., and Gorlin, R. (1976). Problems in echocardiographic volume determinations: echocardiographic-angiographic correlations in the presence of absence of asynergy. Am J Cardiol *37*, 7-11.

Warren, C.R., O'Sullivan, J.F., Friesen, M., Becker, C.E., Zhang, X., Liu, P., Wakabayashi, Y., Morningstar, J.E., Shi, X., Choi, J., *et al.* (2017). Induced Pluripotent Stem Cell Differentiation Enables Functional Validation of GWAS Variants in Metabolic Disease. Cell Stem Cell *20*, 547-557 e547.

Williams, R.R., Rao, D.C., Ellison, R.C., Arnett, D.K., Heiss, G., Oberman, A., Eckfeldt, J.H., Leppert, M.F., Province, M.A., Mockrin, S.C., *et al.* (2000). NHLBI family blood pressure program: methodology and recruitment in the HyperGEN network. Hypertension genetic epidemiology network. Ann Epidemiol *10*, 389-400.

Yu, J., Chau, K.F., Vodyanik, M.A., Jiang, J., and Jiang, Y. (2011). Efficient feeder-free episomal reprogramming with small molecules. PLoS One *6*, e17557.

Yu, J., Hu, K., Smuga-Otto, K., Tian, S., Stewart, R., Slukvin, II, and Thomson, J.A. (2009). Human induced pluripotent stem cells free of vector and transgene sequences. Science *324*, 797-801.

#### SUPPLEMENTAL TABLES

**Table S1. Summary of Generalized Linear Mixed Models (GLMM) and the effects tested†**

| GLMM Model # | Fixed Effects | | | Random Effects | | | | |
|---|---|---|---|---|---|---|---|---|
| | Batch | Post Plate Counts | % Reads Mapped | Experiment Group | Subject | hiPSC Nested in Subject | CMs Nested in hiPSC in Subject | Replicate Nested in CMs in hiPSC in Subject |
| 60 | Y | Y | Y | N | N | N | N | N |
| 61 | Y | Y | Y | Y | N | N | N | N |
| 62 | Y | Y | Y | Y | Y | N | N | N |
| 63 | Y | Y | Y | Y | Y | Y | N | N |
| 64 | Y | Y | Y | Y | Y | Y | Y | N |
| 65 | Y | Y | Y | Y | Y | Y | Y | Y |

**Y**: variable included in the model; **N**: variable not included in the model

†6 models fitted to the gene expression data. All models include 3 fixed effects for experiment batch (Batch), the number of cells that survive the plating process (Post Plate Counts) and percentage of reads successfully mapped (% Reads Mapped). Models #61-#65 each included an additional random effect to account for ET-1 stimulation (Experiment Group), individual donor biological variability (Subject), hiPSC generation noise (hiPSC Nested in Subject), cardiomyocyte differentiation noise (CMs Nested in hiPSC in Subject*)*, and RNA-seq technical noise (Replicate Nested in CMs in hiPSC in Subject).

**Table S2: Variable set enrichment analysis in T genes & GWAS genes of LV mass in HyperGEN African American (AA) samples[††]**

| Pathway/Gene Set Name | Size of Gene set | ∧T | ∧GWAS | FDR for Enrichment in T | FDR for Enrichment in GWAS | T Genes in Pathway | GWAS Genes in Pathway |
|---|---|---|---|---|---|---|---|
| **Overlap with T4** | | | | | | | |
| gross_hypoxia_via_elk3_and_hif1a_up (*Gross et al., 2008; PMID:17704799*) | 142 | 9 | 16 | 0.00009 | 0.00262 | C10orf10 \| EGR1 \| HK2 \| PPAP2B \| RPS6KA5 \| RND1 \| DUSP4 \| ANKRD1 \| PGF | ATF3 \| RORA \| NEDD9 \| MERTK \| ANXA2 \| TNC \| SMOX \| DUSP4 \| MAP3K1 \| ETS1 \| NEDD4L \| CYR61 \| BNIP3 \| NDRG1 \| PALLD \| SLC2A1 |
| mel18_dn.v1_up (Wiederschain et al., 2007; PMID:17452456) | 141 | 6 | 13 | 0.01678 | 0.02855 | KCNN3 \| TIE1 \| FOSL1 \| GPR3 \| CD83 \| CLSPN | KCNN3 \| NRP1 \| WNT5B \| PLAT \| PLAUR \| CHST11 \| NTM \| CDH13 \| F3 \| ERO1L \| LCP1 \| COL13A1 \| WISP1 |
| go_calcium_dependent_phospholipid_binding (*GO:0005544*) | 56 | 4 | 9 | 0.01693 | 0.00487 | MCTP2 \| SYT3 \| SYT10 \| SYTL4 | MCTP2 \| RPH3A \| SYT1 \| DYSF \| PLA2G4A \| ESYT2 \| SYT6 \| ANXA6 \| ANXA2 |
| cahoy_neuronal (*Cahoy et al., 2008; PMID:18171944*) | 100 | 5 | 13 | 0.01894 | 0.00263 | DRD1 \| CPNE4 \| BCL11B \| SH2D5 \| CALN1 | DYNC1I1 \| NELL1 \| TRHDE \| KCNIP4 \| GABRG2 \| MYT1L \| SYT1 \| TSPYL5 \| CNTNAP4 \| CALN1 \| MLIP \| CLSTN2 \| CACNG3 |
| **chen_lvad_support_of_failing_heart_up** (*Chen et al., 2003; PMID:12824457*) | 103 | 5 | 10 | 0.02125 | 0.04555 | ANKRD2 \| ANKRD1 \| C10orf10 \| METTL7A \| AREG | SERPINE1 \| ZBTB16 \| RND3 \| IRS2 \| CYR61 \| ATF3 \| CLU \| CCNB1IP1 \| VCL \| IER5 |
| **weston_vegfa_targets_3hr** (*Weston et al., 2002; PMID:12200464*) | 74 | 4 | 11 | 0.03964 | 0.00260 | PGF \| PIR \| TIE1 \| ANKRD1 | ADD2 \| LMO2 \| BMP6 \| NNMT \| KDR \| RHOB \| CXCR4 \| KRT7 \| ZFHX3 \| EMR1 \| ESR1 |
| corre_multiple_myeloma_up (*Corre et al., 2007; PMID:17344918*) | 74 | 4 | 10 | 0.03964 | 0.00801 | CD200 \| AREG \| ABCA6 \| ANKRD1 | SERPINE1 \| SLC24A3 \| GATA6 \| SAMSN1 \| GLRX \| CLDN11 \| COL13A1 \| RNF145 \| PLAT \| LMO2 |
| **Overlap with T2** | | | | | | | |

**Table S2: Variable set enrichment analysis in T genes & GWAS genes of LV mass in HyperGEN African American (AA) samples[††]**

| Pathway/Gene Set Name | Size of Gene set | ^T | ^GWAS | FDR for Enrichment in T | FDR for Enrichment in GWAS | T Genes in Pathway | GWAS Genes in Pathway |
|---|---|---|---|---|---|---|---|
| nakayama_soft_tissue_tumors_pca1_dn (*Nakayama et al., 2007; PMID:17464315*) | 74 | 6 | 10 | 0.03907 | 0.00801 | COL2A1 \| B3GAT1 \| GPM6B \| SHANK2 \| RIPK4 \| SOX11 | TLE1 \| LHX2 \| ACACB \| PEG3 \| ITIH5 \| GPR125 \| NELL1 \| NPTX2 \| NFIB \| EMX2 |
| servitja_islet_hnf1a_targets_up (*Servitja et al., 2009; PMID:19289501*) | 163 | 8 | 23 | 0.04274 | < 0.00001 | OPLAH \| MAF \| AEBP1 \| CYP1B1 \| CACNA2D3 \| MCAM \| RAB6B \| APOE | KCNIP4 \| ANXA2 \| HMP19 \| MEIS2 \| GAP43 \| PCDH15 \| SNCA \| CDH11 \| NTM \| EPB41L2 \| STXBP5L \| PLA2G4A \| ATF3 \| CACNA2D3 \| STMN2 \| FSTL5 \| EPHA5 \| WISP1 \| DNM1 \| ELAVL2 \| NEBL \| CACNG5 \| AKAP12 |
| **Overlap with T0** | | | | | | | |
| hallmark_epithelial_mesenchymal_transition (*Liberzon et al. 2015; PMID:26771021*) | 200 | 9 | 22 | 0.00251 | 0.00037 | COL11A1 \| QSOX1 \| PCOLCE \| SNTB1 \| MEST \| LEPRE1 \| FMOD \| SNAI2 \| LRP1 | SERPINE1 \| COL4A2 \| NNMT \| THBS1 \| TNC \| CDH11 \| CYR61 \| EDIL3 \| DAB2 \| RHOB \| LEPRE1 \| SGCD \| PLAUR \| MSX1 \| DST \| TPM4 \| GJA1 \| SGCG \| CTHRC1 \| IL15 \| NTM \| SLIT2 |

[††]Only those overlapping gene sets with a size of 200 genes or less are displayed. The number of genes overlapping with the T lists and HyperGEN GWAS results are listed under columns “^T” and “^GWAS”, respectively. Corresponding gene symbols are displayed in the last two columns (7 & 8), separated by “|”.

#### SUPPLEMENTAL FIGURES

**Figure S1. Expression variabilities due to CM differentiation were often smaller than those attributable to hiPSC generation:** This is true in disregard of how variable they might be under ET-1 stimulation. As shown in the barplot, at all significance threshold values tested for differential expression between stimulated and baseline conditions, in a majority of genes, variabilities attributable to CMs cloning are smaller those due to iPSC generation (colored in blue).

**Figure S2. Illustration of the inclusive and exclusive lists of variable (VR) genes:** Both types include 5 lists of gene supposedly sensitive to: ET-1 stimulation (“stimulation”), individual donor biological variability (“individual”), hiPSC generation (“PSC.line”), cardiomyocyte differentiation (“CM.clones”*)*, and RNA-seq technical noise (“technical.noise”). The inclusive “G” lists have non-trivial overlaps (a), while the exclusive “T” lists have no overlap no overlap among them (b).

**Figure S3. Recapture sample clustering:** Hierarchical clustering was applied to expression levels normalized by voom to the residuals of fitting GLMM model #65 in the refined U list of “signal” DE genes. Clones obtained from the same donor were plotted in the same color. Small red circles indicate clones from the only FHM donor.

**Figure S4. Clustering based on GLMM residuals in the U list of “signal” DE genes:** heatmaps of pairwise distances of GLMM residuals over the DE genes: (a) baseline unstimulated cell lines, (b) ET-1 stimulated samples.

**Figure S5. VR genes sensitive to CMs cloning isolate the FHM donor under ET-1 stimulation.** CMs clones made from the only FHM (familial hypertrophic cardiomyopathy) donor were isolated from others using expression of VR genes sensitive to CMs cloning as identified by the quantile method (see Methods). Results from principal coordinate analyses of the expression matrices showed the isolated FHM clone in: (a) unstimulated samples, and (b) ET-1 stimulated samples.

**Figure S6: Effect size of differential target expression of the 4 eQTLs co-localizing with GWAS-detected SNPs:** Regression coefficients from eQTL association analyses in all 43 tissues were extracted from GTEx results and plotted as horizontal bars. Coefficients in LVH relevant tissues were plotted in red. (a) rs309134 on Chr 2 overlap with GWAS of LV mass; (b) rs636049 on Chr 11 also overlap with GWAS of LV mass; (c) rs4244842 on Chr 11 with GWAS of EF; (d) rs11228427 on Chr 11 also with GWAS of EF.

**Figure S1. Expression variabilities due to CM differentiation were often smaller than those attributable to hiPSC generation.**

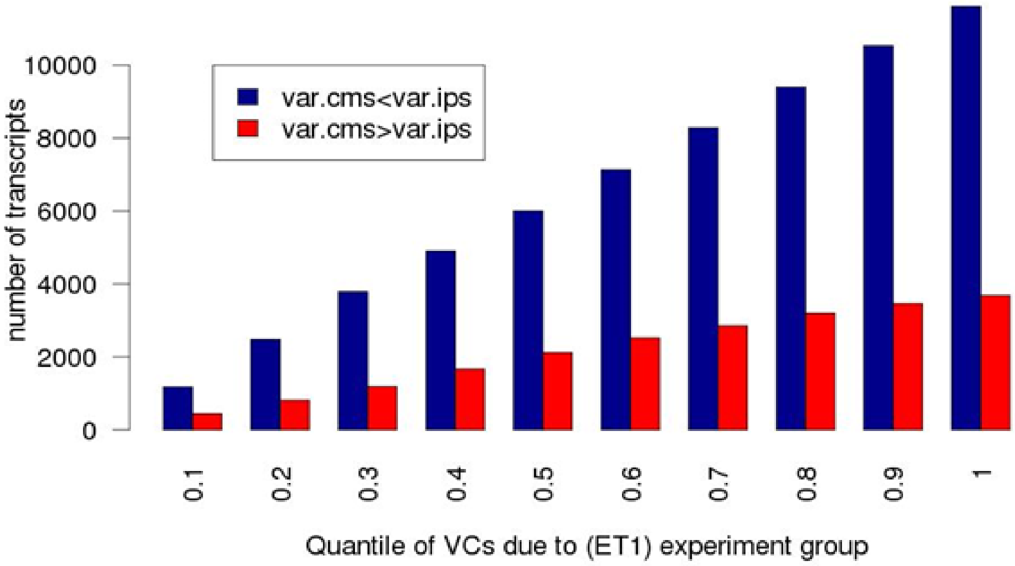

**Figure S2: Illustration of the inclusive and exclusive lists of variable (VR) genes: (a) inclusive G lists (b) exclusive T lists**

(a)

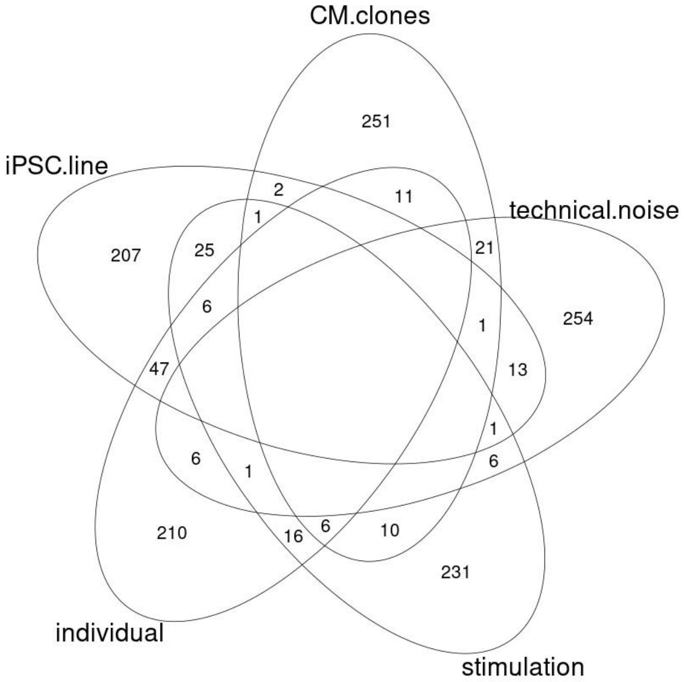

(b)

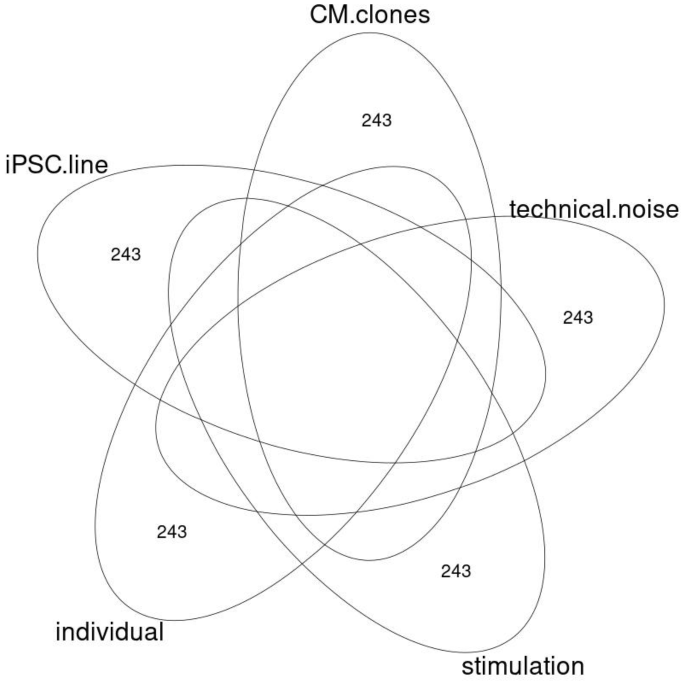

**Figure S3. Recapture sample clustering using GLMM residuals in the refined U list of "signal" DE genes**

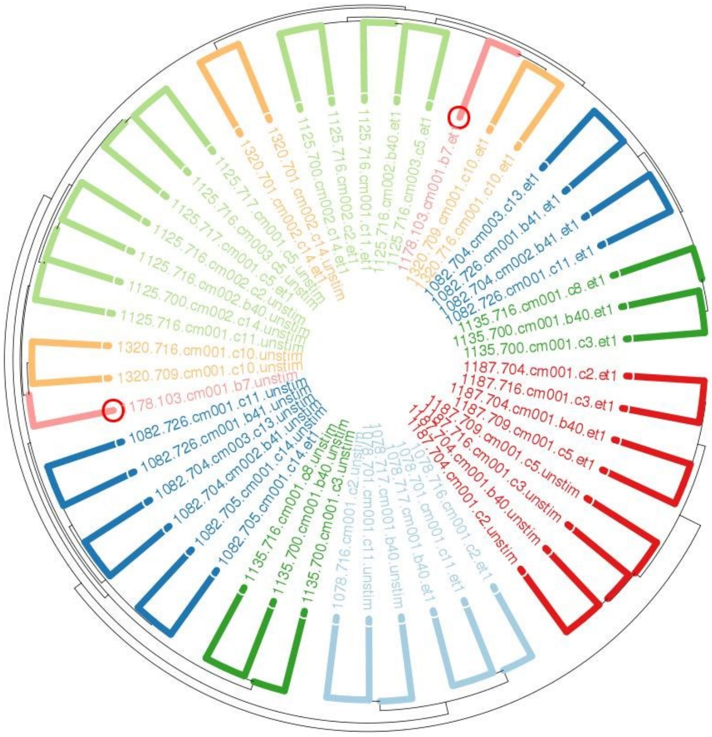

**Figure S4: Clustering based on GLMM residuals in the U list of "signal" DE genes of: (a) unstimulated samples, (b) ET-1 stimulated samples**

(a)

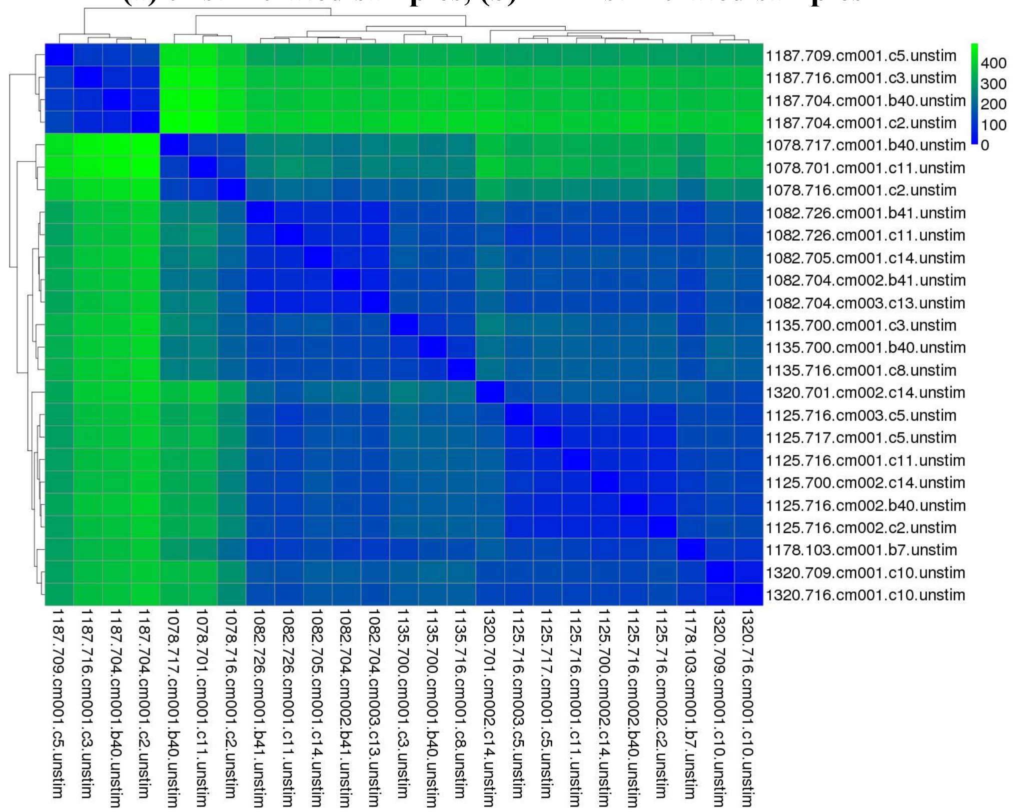

(b)

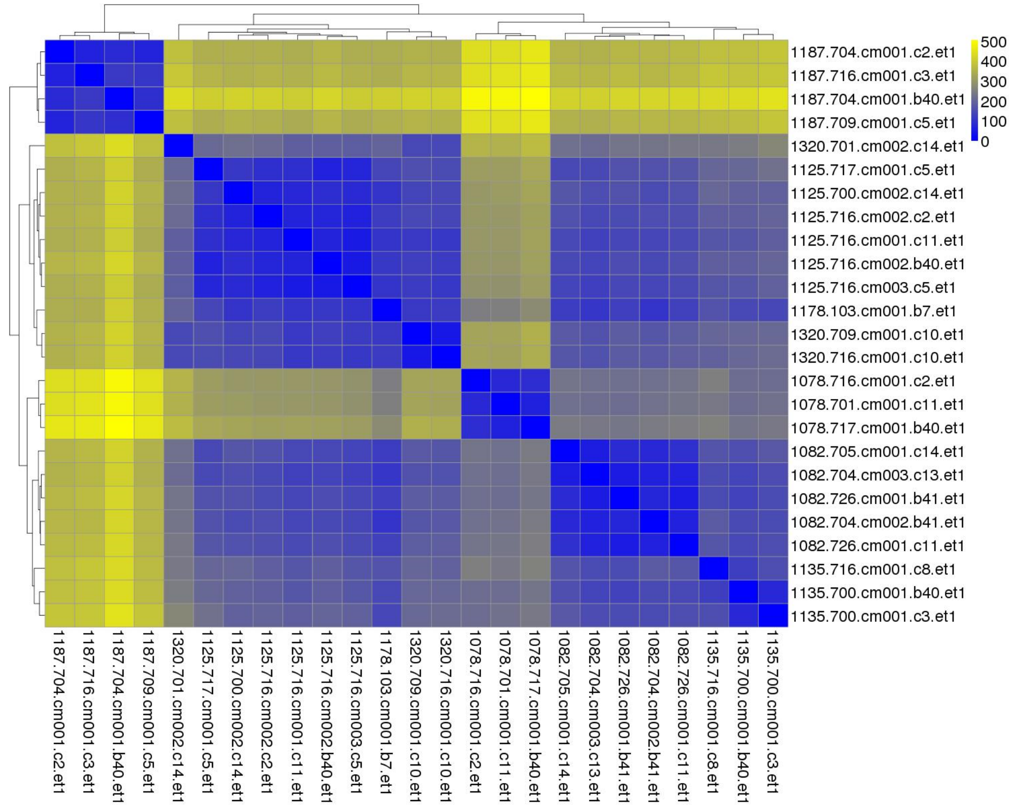

**Figure S5: VR genes due to CMs cloning isolate the FHM donor**

(a)

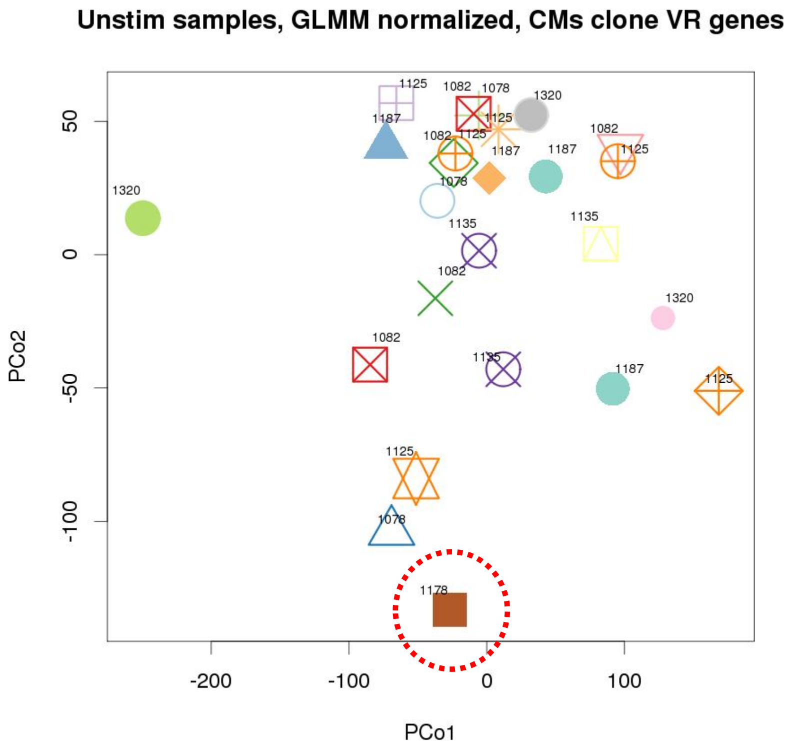

(b)

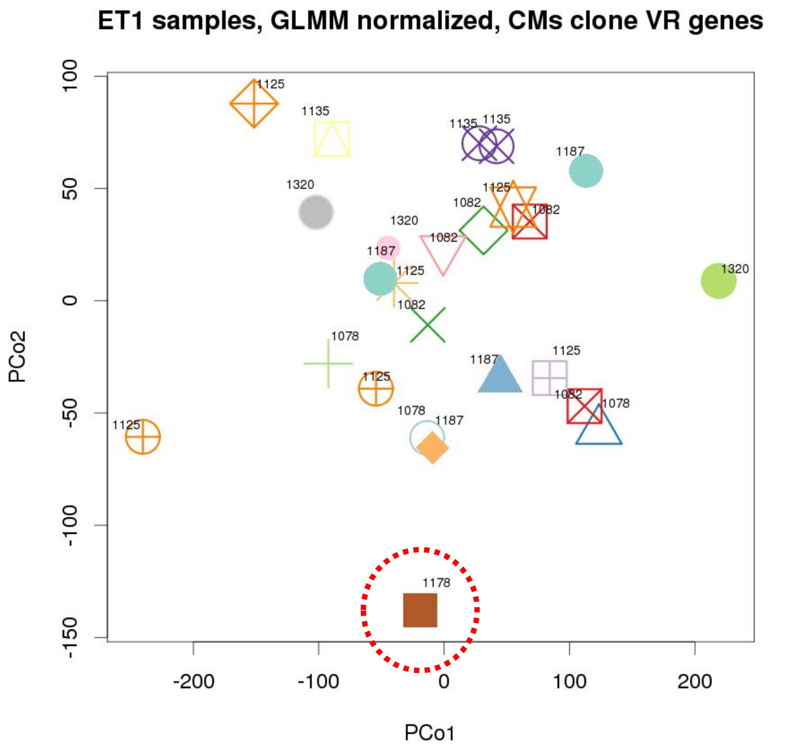

### Figure S6: Effect size of differential expression of the 4 overlapping eQTLs with GWAS

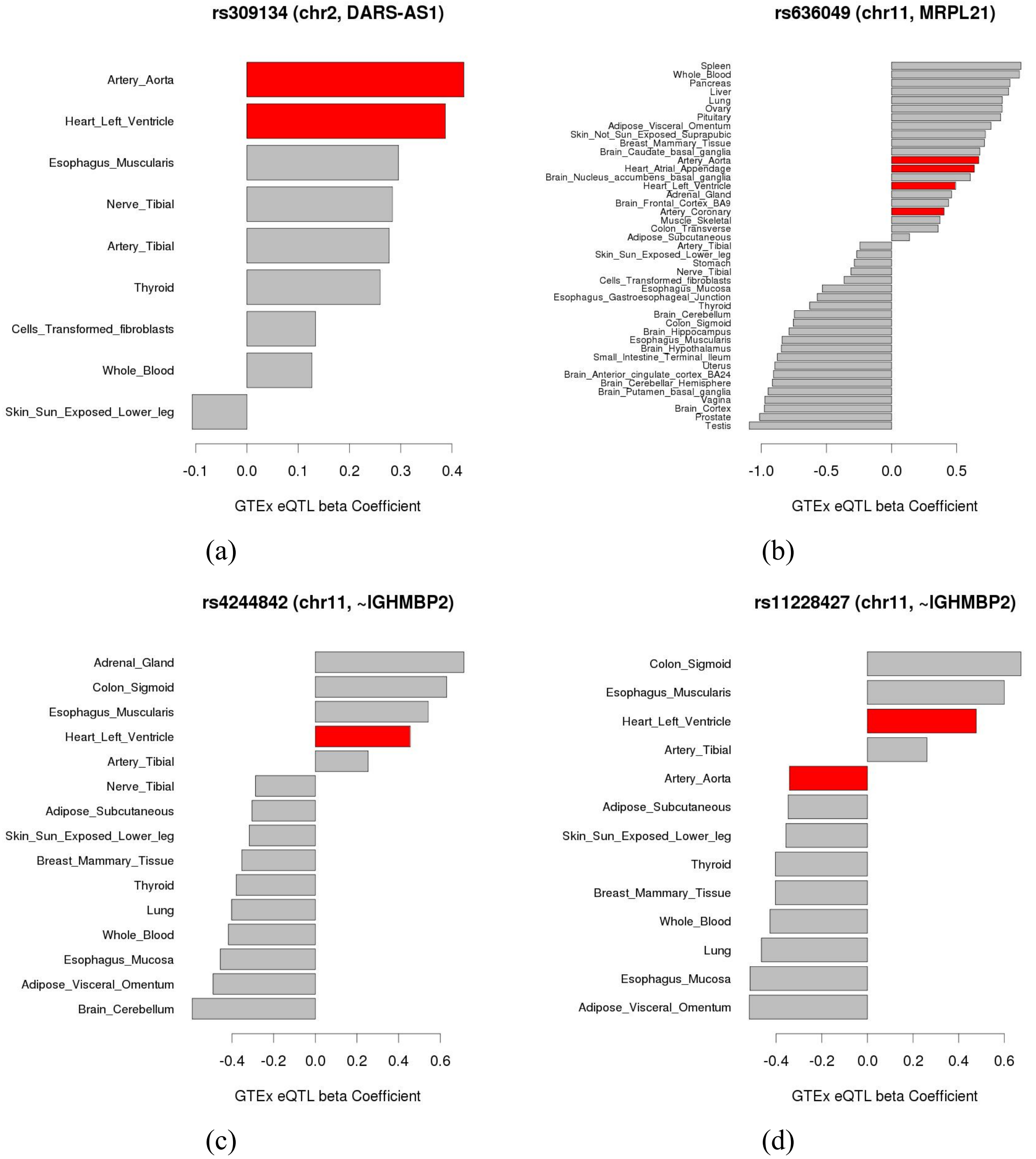

#### Key Resources Table

| REAGENT or RESOURCE | SOURCE | IDENTIFIER |
|---|---|---|
| Antibodies | | |
| Bacterial and Virus Strains | | |
| Biological Samples | | |
| Chemicals, Peptides, and Recombinant Proteins | | |
| Fibronectin (pure) | Sigma-Aldrich | 11051407001 |
| Endothelin-1 | Sigma-Aldrich | E7764 |
| iCell Cardiomyocyte Maintenance Medium | Cellular Dynamics International, Inc | CMM-100-110-005 |
| iCell Cardiomyocyte Plating Medium | Cellular Dynamics International, Inc | CMM-100-110-001 |
| William's E | Thermo Fisher Scientific | A12176-01 |
| Hepatocyte Maintenance Supplement Pack | Thermo Fisher Scientific | CM4000 |
| Critical Commercial Assays | | |
| Total RNA Purification 96-Well Kit | Norgen Biotek Corp. | 24300 |
| Qiagen RNeasy MinElute Cleanup Kit | Qiagen | 74204 |
| Ribo-Zero rRNA Removal Kit | Epicentre | MRZH11124 |
| Ion Torrent Total RNA Seq-Kit v2 | Thermo Fisher Scientific | 4475936 |
| Ion PI™ Template OT2 200 Kit | Thermo Fisher Scientific | A26434 |
| Deposited Data | | |
| Molecular Signatures Database | http://software.broadinstitute.org/gsea/msigdb | N/A |
| Human reference genome NCBI build 37, GRCh37 | http://www.ncbi.nlm.nih.gov/projects/genome/assembly/grc/human/ | |
| Experimental Models: Cell Lines | | |
| HyperGEN-CiPS iPSC lines | Cellular Dynamics International, Inc | |
| HyperGEN-CiPS iPSC-CM lines | Cellular Dynamics International, Inc | |
| FHM cells | Cellular Dynamics International, Inc | |
| Experimental Models: Organisms/Strains | | |
| Oligonucleotides | | |
| Recombinant DNA | | |
| Software and Algorithms | | |
| SAMtools | http://samtools.sourceforge.net/ | Li et al., 2009 |
| R 3.3.0, DESeq2, edgeR, biomRt, reutils, limma, dendextend, circlize, pheatmap | https://www.r-project.org/, https://bioconductor.org/ | |
| RNA-STAR version 2.3.0e | https://github.com/alexdobin/STAR | Dobin et al., 2013 |
| HTSeq version 0.10.0 | https://htseq.readthedocs.io/en/release_0.10.0/ | Anders et al., 2015 |
| Other | | |
| GENCODE v.19 annotation | https://www.gencodegenes.org/releases/19.html | |
